# An Integrated Multi-omics Single Cell Atlas of the Human RPE and Choroid

**DOI:** 10.64898/2026.07.23.740352

**Authors:** Jianming Shao, Xuan Bao, Ismail Yaman, Ye Zheng, Jun Wang, Jin Li, Tingting Yang, Jinjing Jian, Jean Li, Seth Blackshaw, Kapil Bharti, Margo Clarke, Dwight E. Stambolian, Ching-Hwa Sung, Jie J. Zheng, Davide Ortolan, Sarah X. Zhang, Ruchi Sharma, Richard H. Scheuermann, Ajith V. Pankajam, Ahmed H. Ghobashi, Anjun Ma, Qin Ma, Mary Futey, Alexandra-Chloé Villani, J Timothy Stout, Margaret M DeAngelis, Han Chen, Yumei Li, Rui Chen

## Abstract

The retinal pigment epithelium and choroid are critical for supporting the function and maintaining the homeostasis of the outer retina, and their dysfunction underlies a range of inherited and complex ocular diseases. To comprehensively characterize the cellular, transcriptomic, and epigenomic heterogeneity and dynamics within these tissues, we assembled an integrated multi-omics reference atlas comprising 719,813 single-cell/single-nucleus transcriptomes and 234,007 snATAC-seq profiles from 102 ancestrally diverse donors spanning 0 to 99 years of age, including cells from both the macula and periphery. This atlas resolves 48 distinct cell types or states and catalogs 448,567 open chromatin regions. Specifically, we resolved five distinct RPE subpopulations organized along a central-to-peripheral spatial axis, alongside two distinct stress/senescence states. We reconstructed the transcriptomic and epigenetic zonation of endothelial cells and expanded choroidal stromal heterogeneity by characterizing 11 fibroblast and two pericyte types. Age-associated compositional analysis revealed a significant fractional depletion of melanocytes, PI16+ fibroblasts, and venule endothelial cells with age, alongside a modest relative loss of central RPE and a corresponding increase in far-peripheral RPE. Cell-type-specific aging transcriptomics uncovered shared pathways related to inflammatory responses alongside distinct cell-type-specific signatures. Notably, significant age-associated epigenetic changes concentrated in the macula during the transition from early-to-middle adulthood and remained stable into old age, with transcription factors from the AP-1/bZIP family emerging as the dominant enriched motifs. Finally, integrating this atlas with AMD GWAS data provides novel variant-to-gene evidence implicating *LIPG* and *COL4A3* in AMD pathogenesis. Together, this multi-omics atlas serves as both an invaluable community reference and a powerful discovery engine that translates genetic risk signals into localized target cells and candidate mechanisms, laying a foundation for understanding RPE/choroid biology in health and disease.

## Introduction

The human eye is a highly specialized organ designed to capture, focus, and transduce light into neural signals. The neural retina plays a critical role in this process, serving as the primary site for the initial transduction and processing of visual information. The health and function of these retinal circuits, particularly the photoreceptors, depend on the support of the retinal pigment epithelium (RPE)—a specialized monolayer of pigmented cells critical for visual cycle maintenance, nutrient transport, immune privilege, and the phagocytosis of shed photoreceptor outer segments^1^. Supporting the RPE and outer retina is the choroid, a highly vascularized tissue that provides the primary blood supply to the outer retina while facilitating thermal regulation and waste removal^2^. Dysfunction within the RPE-choroid complex is a primary driver for a spectrum of inherited and complex ocular pathologies, most notably age-related macular degeneration (AMD).

To better characterize the pathogenesis of ocular diseases, it is essential to establish a comprehensive reference cell map to better understand cellular heterogeneity, composition, and the dynamic transcriptomic and epigenetic changes that occur across demographic variables in the retina, RPE, and choroid. Established in 2016, the Human Cell Atlas (HCA) consortium aimed to create a comprehensive biological map of all cells within the human body^3,4^. To date, it has released several tissue-specific atlases, including the brain^5^, lung^6^, breast^7^, heart^8^, liver^9^ and immune system^10^. As a core component of the HCA, the Eye Biological Network aims to build a comprehensive molecular and spatial atlas of the visual system; it has already released version 1 of the single-cell atlas for the human retina^11^.

Recognizing the vital roles of the RPE/choroid, several recent single-cell and single-nucleus omics studies have begun to establish “cell atlases” of these tissues^12–24^. These atlases provide a transformative framework for dissecting cellular complexity, enabling the mapping of specific regulatory landscapes in both healthy and diseased eyes. However, existing studies remain constrained by several limitations: (1) limited number of cells profiled, particularly for the RPE, which remains insufficient to move beyond broad cell-class annotations; (2) under-characterized heterogeneity within major cell classes; and (3) incomplete aging trajectories due to a lack of large-scale, age-diverse cohorts.

To address these limitations, we present an in-depth, comprehensive human RPE/choroid cell atlas. This integrated resource comprises 719,813 cells derived from 154 samples across 102 donors, encompassing a broad age distribution and diverse ancestral backgrounds, including 551,214 snRNA-seq and 168,599 scRNA-seq profiles. This atlas enabled the identification of 17 major cell classes and 48 subtypes or states, including five distinct RPE subpopulations. Furthermore, we identified 448,567 Open Chromatin Regions (OCRs); an integrative analysis with snRNA-seq data allowed us to infer the gene expression associated with OCR accessibility. Finally, we leveraged single-cell spatial transcriptomics to resolve the structural architecture and spatial organization of the RPE/choroid complex. Using this multi-omic framework, we explored the cellular composition, transcriptomic, and epigenetic dynamics across the human lifespan. To demonstrate the utility of our dataset, we performed functional prioritization of AMD GWAS variants and proposed novel therapeutic targets. Collectively, our comprehensive RPE/choroid multi-omics resource lays a foundation for investigating the mechanisms of, and developing treatments for, inherited and complex ocular diseases.

## Results

### Single-Cell and Single-Nucleus Transcriptomic Atlas of the Human RPE/Choroid

To construct a comprehensive cell atlas of the human retinal RPE and choroid, we integrated nine publicly available single-cell and single-nucleus RNA-seq (sc/snRNA-seq) datasets^13,15,16,18,19,22,23^ with newly generated data. Our new contributions account for 23.83% of the total scRNA-seq and 92.69% of the total snRNA-seq cells in the final resource (Figure 1a, Table S1). This integrated atlas comprises 719,813 cells—consisting of 551,214 snRNA-seq and 168,599 scRNA-seq profiles—derived from 154 samples across 102 donors with a broad age distribution (0–99 years) (Figure 1b, Table S1). To ensure demographic consistency, we performed germline variant calling from the scRNA-seq data to infer genetic ancestry for 54 public donors lacking this information and cross-validated demographic metadata for our newly generated cohort of 48 donors (Table S1). Our 48-donor cohort encompasses diverse ethnicities (25% European, 33.33% African, and 35.42% Hispanic, and 6.25% Asian) and a balanced sex distribution (45.28% female vs. 54.72% male) (Figure 1c, Table S1). The atlas incorporates cells from both the macular and peripheral regions of the posterior segment. While our newly generated data were derived from healthy donors, the integrated atlas includes 58,210 AMD-associated cells from the incorporated public datasets (Table S1).

**Figure 1.**
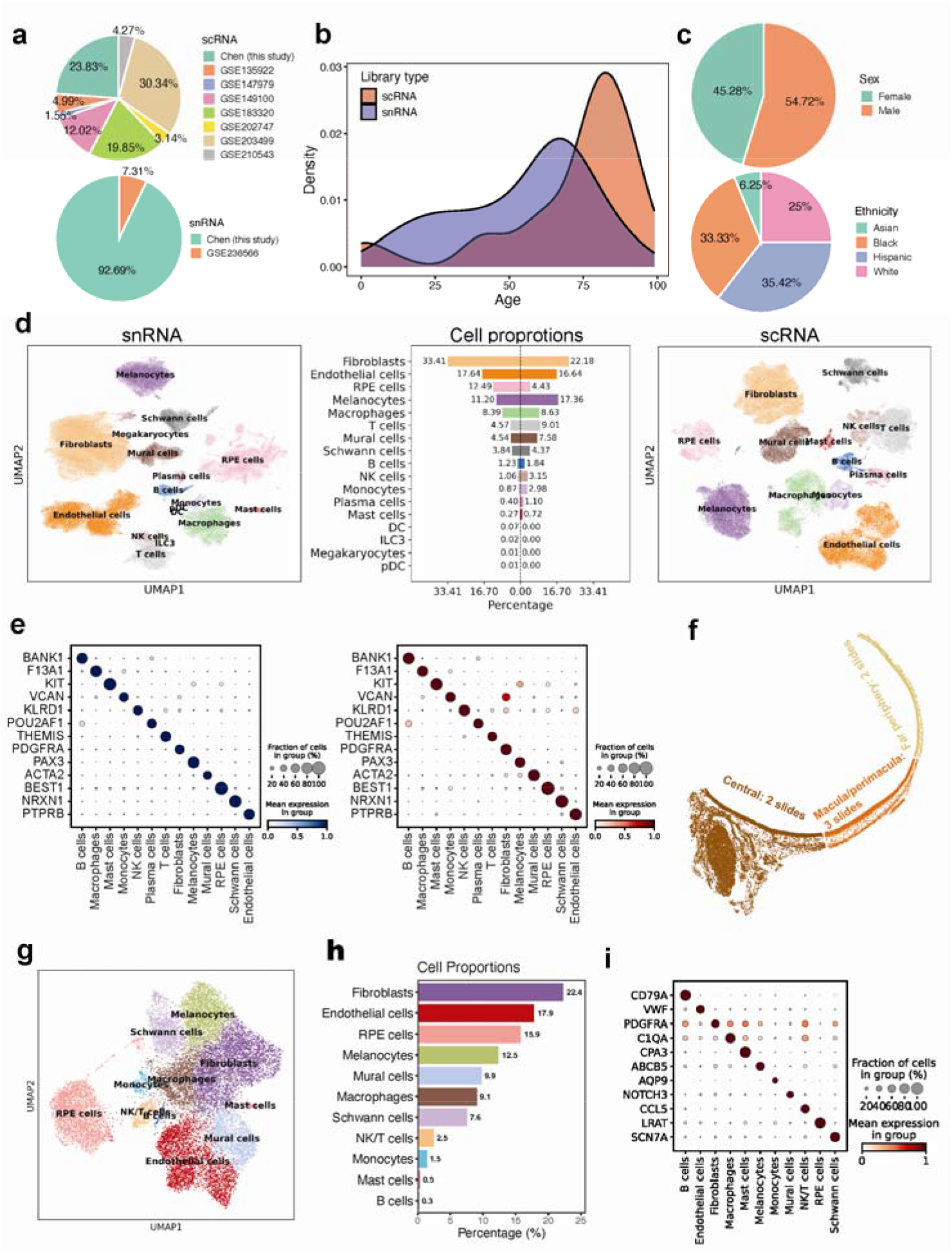
Integrated single-cell, single-nucleus, and spatial transcriptomic landscape of the human RPE/choroid. **a,** Pie charts summarizing the contribution of cells from the present study and previously published datasets to the integrated atlas, shown separately for scRNA-seq (top) and snRNA-seq (bottom). **b,** Density plots illustrating the age distribution of donors for the scRNA-seq and snRNA-seq cohorts. **c,** Pie charts depicting the sex (top) and genetic ancestry (bottom) distributions of donors within the snRNA-seq dataset. **d,** UMAP visualizations of major cell classes for the snRNA-seq (left) and scRNA-seq (right) modalities. Middle: A bidirectional bar plot comparing the relative proportions of major cell classes between modalities. **e,** Dot plots showing the expression profiles of canonical marker genes across major cell classes in snRNA-seq (left) and scRNA-seq (right) data. **f,** Schematic representation of the posterior segment regions profiled using the Xenium spatial platform. **g,** UMAP visualization of major cell classes identified in the Xenium spatial transcriptomics data. **h,** Bar plot illustrating the cellular composition of the Xenium dataset. **I,** Expression profiles of marker genes across major cell classes within the Xenium spatial data.

Using scVI^25^, we performed multi-sample batch correction and dataset integration, processing the scRNA-seq and snRNA-seq modalities separately. Both modalities successfully captured all major reported cell types of the RPE/choroid complex—including fibroblasts, endothelial cells, RPE cells, melanocytes, macrophages, T cells, mural cells, Schwann cells, B cells, NK cells, monocytes, plasma cells, and mast cells—as confirmed by the expression of established lineage-specific marker genes (Figures 1d–e). Rare immune populations, such as dendritic cells (DCs), ILC3s, pDCs, and megakaryocytes, were identified in the snRNA-seq dataset but were absent in the scRNA-seq data, likely due to the smaller total number of cells profiled in the latter modality (Figure 1d). Each major cluster contained cells from multiple donors with comparable sequencing quality (Figure S1a, Table S2). Cross-comparison of our refined annotations with the original cell labels from Monavarfeshani et al.^23^ confirmed high consistency, further validating our cell-type assignments within the integrated atlas (Figure S1b). Overall cellular compositions were consistent across modalities, with fibroblasts, endothelial cells, and melanocytes identified as the most abundant populations (Figure 1d). Notably, the snRNA-seq component of the atlas captured 68,871 RPE cells, a significant increase compared to the 7,472 cells identified in the combined scRNA-seq datasets (Figure 1d). Finally, high-resolution sub-clustering of the snRNA-seq data (n = 551,214 cells) yielded 48 distinct cellular subtypes and states (Figure S1c).

To evaluate the transcriptomic similarities and inherent biases between the two modalities, we systematically compared the profiles of scRNA-seq and snRNA-seq across all major cell types. NS-Forest^26^ identified shared core transcriptional signatures, with the top binary markers exhibiting high consistency between modalities (Table S3). Cross-modality co-embedding using sysVI^27^ and transcriptomic similarity analysis via MetaNeighbor^28^ confirmed the robust alignment of corresponding cell types (Figures S2a, S2b). Differential detection gene (DDG) analysis identified significant modality-specific signatures, with an average of 6,438 DDGs per major cell class (range: 2,518–11,653; Figure S2c, Table S4). While the majority of DDGs were shared by at least two cell types, 27.01% were unique to a specific cell type. Gene Ontology (GO) enrichment analysis revealed that genes preferentially detected in scRNA-seq were enriched for processes including translation, mRNA processing, and ribosome biogenesis (Figure S2d). Conversely, snRNA-seq-enriched genes were associated with small GTPase-mediated signal transduction, protein modification, and organelle organization (Figure S2e).

We performed Xenium spatial transcriptomics profiling of the temporal posterior segment using a custom gene panel (Table S5). A total of seven slides—encompassing the central posterior segment (n=2), partial macula and periphery (n=3), and far periphery (n=2)—were utilized to map the spatial distribution of the identified cell types (Figure 1f). After filtering for neural retina and optic nerve cells, we recovered 18,157 segmented cells (Figure 1g). The Xenium data successfully captured 11 of the major cell types identified in the snRNA-seq atlas, exhibiting consistent cellular compositions (Figures 1g–h) and spatially restricted expression of canonical marker genes (Figure 1i). The spatial transcriptomic data co-embedded robustly with both the snRNA-seq and scRNA-seq datasets, though we observed a slightly higher integration concordance with the scRNA-seq modality (Figure S3). These findings validate the high quality of the Xenium spatial data.

### Single-Nucleus Chromatin Accessibility Atlas of the Human RPE/Choroid

While transcriptomic profiling defines cell identity and composition, the underlying epigenetic landscape governs the regulatory programs that maintain these states. To define the cis-regulatory architecture of the RPE/choroid, we generated snATAC-seq chromatin accessibility profiles for 234,007 nuclei across 55 samples from 41 donors, recovering 92,001 macular and 142,006 peripheral nuclei (Table S6). The cohort features diverse ancestral backgrounds (34.15% European, 36.59% African, and 26.83% Hispanic, 2.43% Asian), a balanced sex distribution (43.90% Female vs. 56.10% Male), and a broad age range (0–93 years) (Table S5). Cross-modal integration, using snRNA-seq labels as a reference, captured 12 major cell types (Figure 2a); certain rare immune populations were likely missed due to the relatively small number of cells profiled in this modality. Each major cluster comprised cells from multiple donors with comparable sequencing quality and consistent multi-donor contributions (Table S7). Cellular compositions were concordant with the snRNA-seq atlas (Figure 2b), and canonical marker genes exhibited strong correlation between promoter chromatin accessibility and gene expression (Figure 2c).

**Figure 2.**
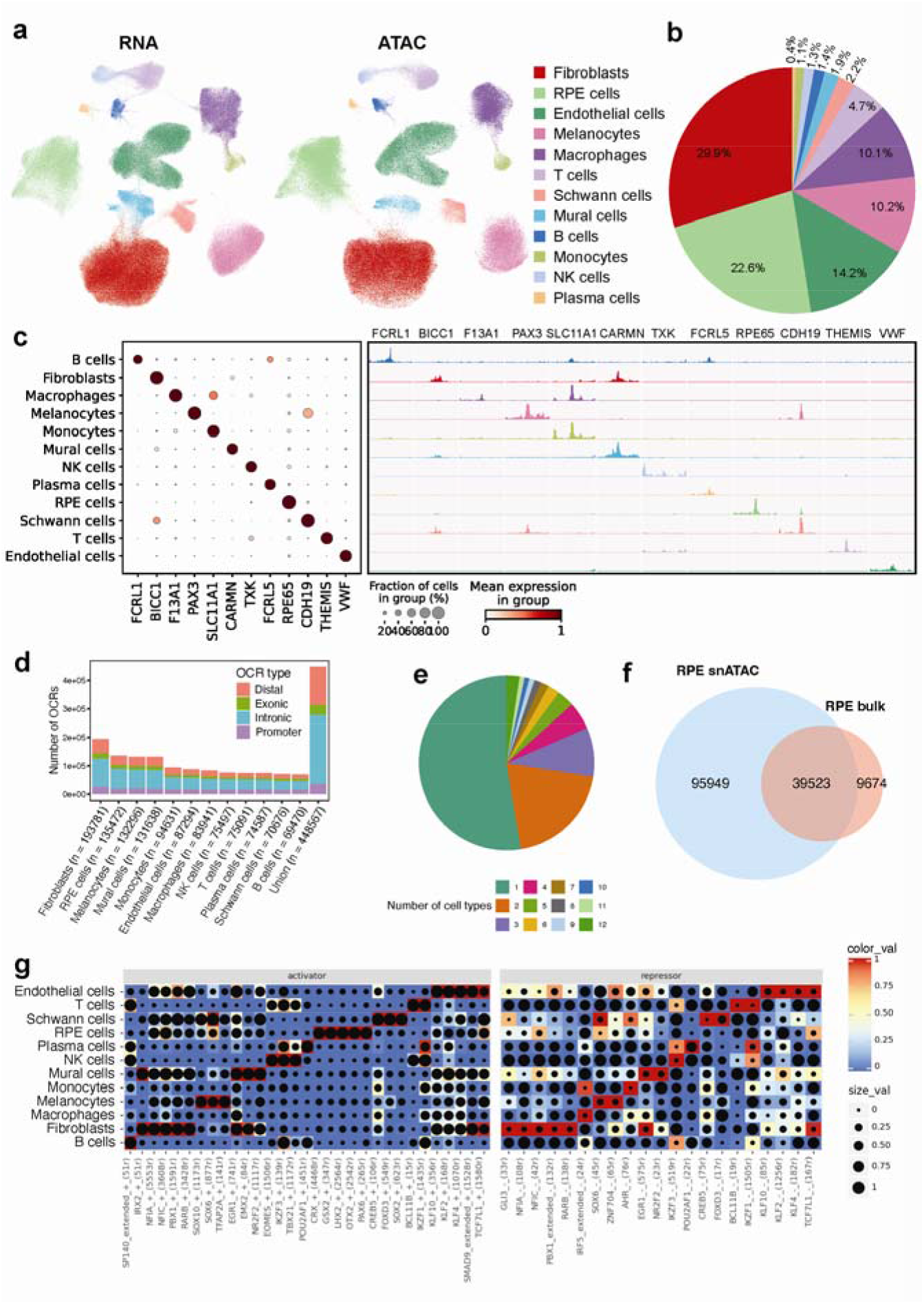
Single-nucleus chromatin accessibility landscape of the human RPE/choroid. **a,** UMAP visualization of integrated snRNA-seq and snATAC-seq data, demonstrating consistent co-embedding and successful cross-modal alignment of major cell classes. **b,** Pie chart showing the proportional distribution of major cell classes identified within the RPE/choroid snATAC-seq atlas. **c,** Integrated view of marker gene expression and accessibility. Left: Dot plot showing marker gene expression across cell classes in snRNA-seq. Right: Pseudo-bulk accessibility tracks (snATAC-seq) at the promoter regions of corresponding markers. **d,** Bar plot illustrating the total number of open chromatin regions (OCRs) identified per cell class, categorized by genomic feature (distal, exonic, intronic, or promoter). **e,** Pie chart summarizing the cell-type specificity of OCRs, indicating the number of cell classes in which each peak is detected. **f,** Venn diagram illustrating the overlap of RPE derived OCRs identified in the current snATAC-seq dataset compared with a published bulk RPE ATAC-seq study by Wang et al. **g,** Heatmap of core regulons identified by SCENIC+, displaying both activator (left) and repressor (right) motifs. Dot size indicates TF motif enrichment within cell-type-specific peaks, while color intensity represents the integrated regulon activity score.

We identified 448,567 open chromatin regions (OCRs), ranging from 69,470 to 193,781 per cell type (Figure 2d), with 52.7% exhibiting cell-type-specific accessibility (Figure 2e). Notably, our snATAC-seq data captured 80.34% of OCRs reported in existing bulk RPE ATAC-seq studies while expanding the total number of discovered OCRs nearly three-fold (Figure 2f). Differential accessibility analysis identified between 492 and 11,497 differentially accessible regions (DARs) per cell class (FDR ≤ 0.01, log2FC ≥ 1; Figure S4a, Table S8). Motif enrichment of these DARs predicted transcription factor (TF) activities consistent with established literature, such as OTX2 in RPE^29^, SOX4/MITF in melanocytes^30,31^, and SPI1/SPIC in macrophages^32,33^ (Figure S4b). Co-accessibility analysis identified 143,823 distal peaks correlated with proximal promoters (score > 0.3), defining 294,601 predicted enhancer-promoter interactions (Figure S4; Table S9). Integration of snATAC-seq and scRNA-seq further identified 183,327 OCRs as putative cis-regulatory elements (CREs) across 420,724 peak-gene pairs (correlation > 0.3, FDR < 0.01; Figure S5; Table S10). Finally, SCENIC+ integration of the snRNA and snATAC modalities inferred core eRegulons for the major cell types, identifying TFs that were also detected in our DAR motif enrichment, including key regulators of RPE and choroidal cell specification (Figure 2g, Figure S5).

### Regional Divergence in Cellular Composition, Transcriptomics, and Chromatin Accessibility Across the Macula and Periphery

Given the substantial number of cells captured from both macular and peripheral regions across a broad age distribution, we first investigated age-related shifts in relative cellular composition. Analysis of these trajectories revealed significant increases in peripheral mural cells (R = 0.37, P = 0.011) and Schwann cells (R = 0.38, P = 0.0093), alongside a decrease in peripheral melanocytes (R = -0.39, P = 0.0075); a similar downward trend was observed for macular melanocytes, though it did not reach statistical significance (R = -0.34, P = 0.10) (Figure 3a). Endothelial cells (ECs) displayed divergent regional trajectories: peripheral EC proportions trended positively with age (R = 0.24, P = 0.095), while macular EC proportions exhibited a negative trend (R = -0.25, P = 0.25) (Figure S6a). In contrast, fibroblast and RPE proportions remained stable across the lifespan. Within immune populations, we noted a marginal increase in peripheral plasma cells (R = 0.43, P = 0.054), a decrease in macular monocytes (R = -0.41, P = 0.068, Figure S6b), and a significant increase in macular T cells (R = 0.42, P = 0.05; Figure S6c).

**Figure 3.**
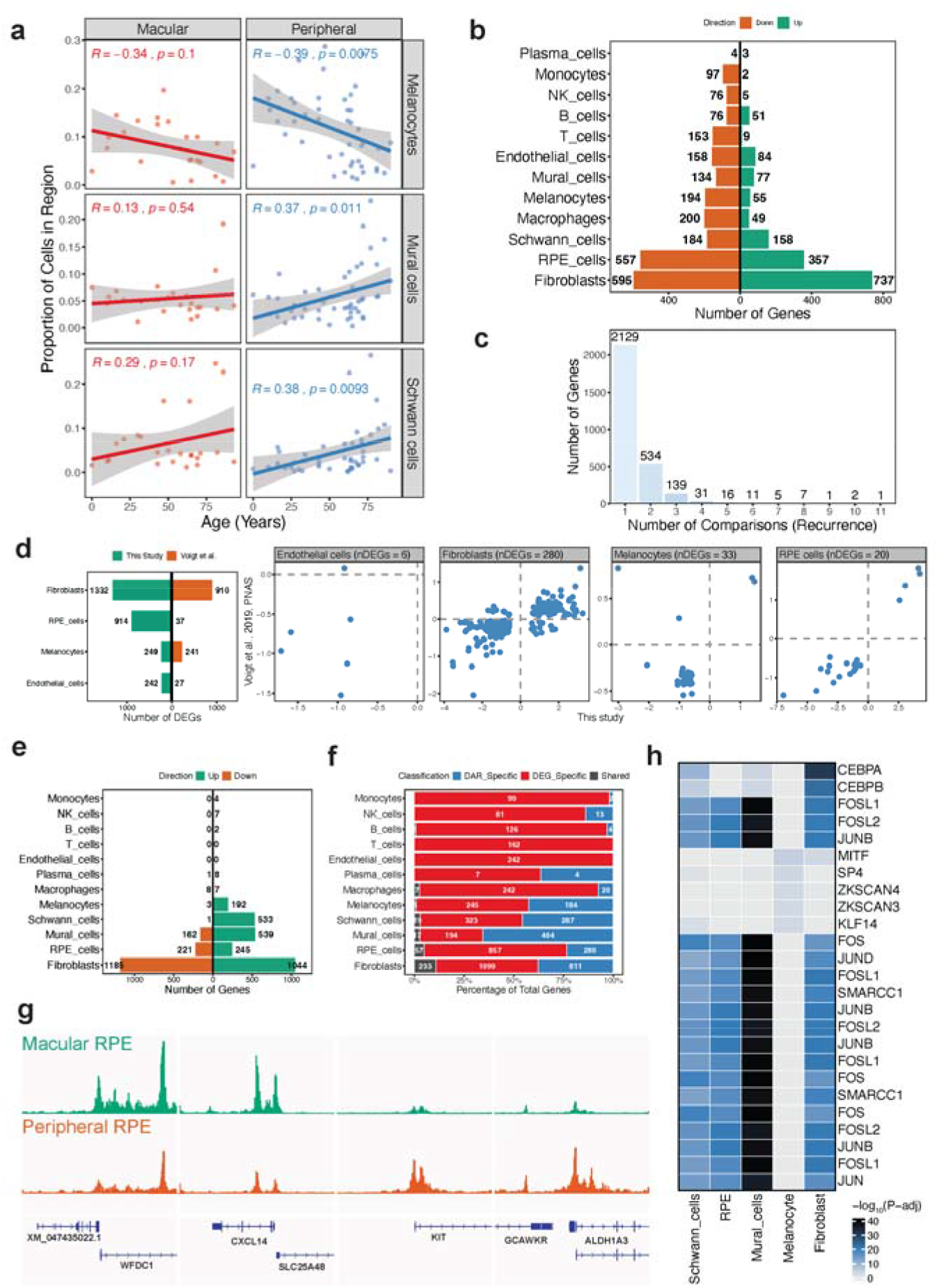
Regional Divergence in Cellular Composition, Transcriptomics, and Chromatin Accessibility Across the Macula and Periphery. **a,** Age-associated shifts in cellular composition for macular and peripheral melanocytes, mural cells, and Schwann cells. **b,** Regional transcriptomic divergence shown by a bi-directional bar plot indicating the number of macular-enriched and peripheral-enriched DEGs across 12 major cell types. **c,** Sharing of regional DEGs across cell types, highlighting genes specific to or shared among various lineages. **d,** Comparative analysis of regional DEGs identified in this study versus those reported by Voigt et al. **e,** Regional epigenetic divergence shown by a bi-directional bar plot indicating the number of macular-high and peripheral-high DARs across 12 major cell types. **f,** Integration of transcriptomic and epigenetic signatures via cross-referencing regional DEGs with regional DAR-associated genes. **g,** Representative genome tracks visualizing differential chromatin accessibility at the promoters of RPE regional markers, including macular-enriched (*WFDC1*, *CXCL14*) and peripheral-enriched (*KIT*, *ALDH1A3*) genes. **h,** Transcription factor (TF) motif enrichment showing the top 5 enriched motifs in macular-high DARs across five cell types with sufficient DAR counts for robust analysis.

We subsequently investigated regional differentially expressed genes (DEGs), specifically focusing on donors with matched samples from both macular and peripheral regions. This analysis identified between 9 and 1,336 regional DEGs across 12 cell types (FDR ≤ 0.05, |FC| ≥ 1.5; Figure 3b, Table S11). Fibroblasts exhibited the highest regional variation (736 macular-upregulated, 600 macular-downregulated), followed by RPE cells (911 DEGs), whereas immune cells exhibited fewer regional differences. While most DEGs were cell-type specific, 735 genes were shared by at least two cell types (Figure 3c, Figure S7a). Comparisons with a prior study^13^ demonstrated high directional consistency, with all 20 shared RPE DEGs regulated in the same direction (Figure 3d, Table S12). Pathway analysis indicated that peripheral-upregulated genes were enriched for epithelial-mesenchymal transition (EMT), hypoxia, UV response, TNF-alpha/NF-kB, and IL-2/STAT5 signaling, while macular-upregulated genes showed significant pathway enrichment only in fibroblasts (Figure S7b). Notably, the intersection of these DEGs with AMD risk genes identified regionally biased expression patterns, including *VEGFA*, *TIMP3*, *CFB*, *COL4A3*, *HTRA1*, and *ABCA1* in fibroblasts; *ABCA1* and *COL8A1* in RPE; *C3* and *TRPM3* in macrophages; and *TSPAN10* and *TIMP3* in melanocytes (Figure S7c).

To determine if these transcriptomic differences are underpinned by epigenetic shifts, we performed differential accessibility analysis across all identified cell types. While almost no differentially accessible regions (DARs) are observed in immune cell and endothelial cells (Figure 3e, Figure S8, Table S13), a large number of DARs are found in melanocytes, Schwann cells, and mural cells. A higher number of DARs with increased accessibility in the macula for mural cells, while RPE and fibroblasts showed more balanced distributions (Figure 3e). By linking DARs to their putative target genes, we identified between 2 and 1,044 DAR-associated genes per cell type (Figure 3f). In RPE and fibroblasts, we observed a modest correlation between DAR-associated genes and regional DEGs; for example, macular RPE markers *CXCL14* and *WFDC1* exhibited increased macular chromatin accessibility, while peripheral markers *KIT* and *ALDH1A3* showed higher peripheral accessibility (Figure 3g). Motif analysis of macular-enriched DARs identified an enrichment for *MITF*, *SP4*, *ZKSCAN3/4*, and *KLF14* in melanocytes, and AP-1 family members (FOSL1, FOSL2, FOS, JUN, JUNB) across the remaining four non-immune cell types (Figure 3h).

### Transcriptomic and spatiotemporal heterogeneity of RPE cells

In this study, we recovered a total of 68,871 high-quality RPE nuclei, comprising 10,568 from the macula and 58,303 from the periphery. Re-clustering of these nuclei identified five clusters with distinct transcriptomic profiles (Figure 4a), each exhibiting high sequencing quality and contributions from multiple donors (Figure S9a, Table S14). Cluster-specific marker genes were identified using NS-Forest (Figure 4b, Table S15). Notably, the *TUBA1A*+ RPE cluster contained a significantly higher proportion of cells from aged donors (Figure 4c). Regional distributions also varied significantly: RPE *LGI1*+ contained the highest proportion of macular cells, followed by RPE *TUBA1A*+ and RPE *CST3*+ (Figure 4c). Pseudobulk analysis identified an average of 1,048 cluster-specific DEGs (Table S16, Figure S9b). Pathway analysis revealed that RPE *CST3*+ and RPE *TUBA1A*+ were enriched for the unfolded protein response (UPR), inflammatory signaling, hypoxia, and DNA repair (Figure 9c).

**Figure 4.**
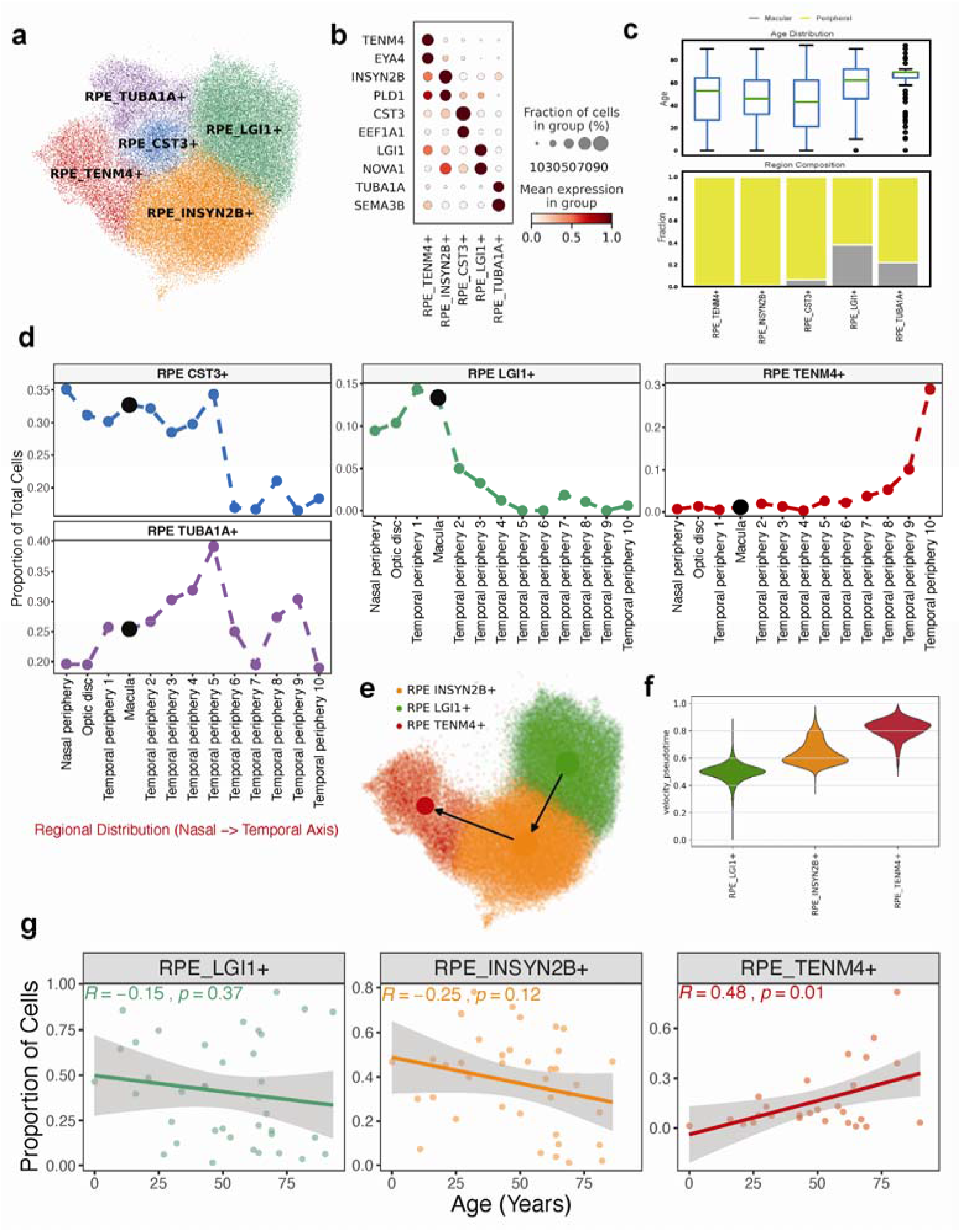
Transcriptomic and spatiotemporal heterogeneity of RPE cells. **a,** UMAP visualization of 68,871 high-quality RPE nuclei identifies five distinct transcriptomic clusters: *LGI1*+, *CST3*+, *INSYN2B*+, *TENM4*+, and *TUBA1A*+. **b,** Dot plot illustrating the expression of top marker genes defining each RPE type. Dot size indicates the percentage of cells expressing the gene, and color intensity represents the mean expression level. **c,** Box plots (top) showing the age distribution of donors contributing to each cluster, and a stacked bar plot (bottom) showing the relative proportion of macular versus peripheral cells within each cluster. **d,** Regional distribution across a nasal-to-temporal axis (spanning from the nasal periphery to temporal periphery 10) for *CST3*+, *LGI1*+, *TENM4*+, and *TUBA1A*+ clusters. **e,** PAGA velocity graph illustrating a predicted spatial continuum and transcriptomic transition originating from RPE *LGI1*+, progressing through *INSYN2B*+, and terminating at *TENM4*+. **f,** Violin plot displaying the distribution of velocity pseudotime across the three spatial clusters. **g,** Age-associated shifts in cellular composition for RPE *LGI1*+, RPE *INSYN2B*+ and RPE *TENM4*+.

Based on the distinct regional distributions of these clusters, we investigated a potential spatial axis extending from the macular center to the periphery shows a transition from RPE *LGI1*+ to *INSYN2B*+ and finally to *TENM4*+ populations. To validate this spatial organization and further improve the resolution, we performed marker-based annotation of the RPE within our Xenium spatial transcriptomics dataset (Figure S10a, 10b). Although limited panel representation precluded the identification of the RPE *INSYN2B*+ cluster, our spatial data successfully resolved the distinct localization patterns of the remaining four RPE populations (Figure 4d, Figure S10b). Spatial transcriptomics mapped *LGI1*+ RPE around the macular, while *TENM4*+ RPE was localized to the far periphery (Figure 4d, Figure S10a). RPE *CST3*+ cells showed a central-to-mid-peripheral distribution, while *TUBA1A*+ RPE cells was observed across all regions.

There is limited panel representation that precluded the identification of the RPE *INSYN2B*+ cluster. To infer the localization of the RPE *INSYN2B*+ population, we utilized RNA velocity to model cellular transitions. PAGA velocity graphs revealed a directional trajectory transitioning from the *LGI1*+ cluster, through the *INSYN2B*+ state, and culminating in the *TENM4*+ population (Figure 4e-f). Pseudotime-associated gene analysis (Monocle3) identified 1,031 genes correlated with the peripheral transition (enriched for hypoxia and TGF-beta signaling) and 480 genes correlated with central localization (enriched for cholesterol homeostasis and angiogenesis) (Table S17, Figure S10b-d). Finally, age-related composition analysis revealed a significant increase in the RPE *TENM4*+ proportion with age (R = 0.48, P = 0.01), alongside positive trends for RPE *CST3*+ and *TUBA1A*+, and negative trends for RPE *LGI1*+ and *INSYN2B*+ (Figure 4g, Figure S10e).

### Dynamics of Cellular Senescence, Transcriptome, and Chromatin Accessibility in the Aging RPE

We next investigated the shifts in cellular senescence, transcriptomic profiles, and chromatin accessibility occurring in the RPE during aging. We first characterized RPE senescence by inferring cellular senescence states (Methods). The proportion of inferred senescent RPE cells correlated positively with donor age (R = 0.41, P = 0.029; Figure 5a) and was significantly higher in the macula than in the periphery (P = 0.025, Figure 5b). Furthermore, senescent cells were enriched within the *CST3*+ and *TUBA1A*+ clusters (Figure 5c). Enrichment analysis of senescence-associated genes highlighted pathways involved in the EMT, IL-6/JAK/STAT3 signaling, inflammatory responses, and apoptosis (Figure 5d).

**Figure 5.**
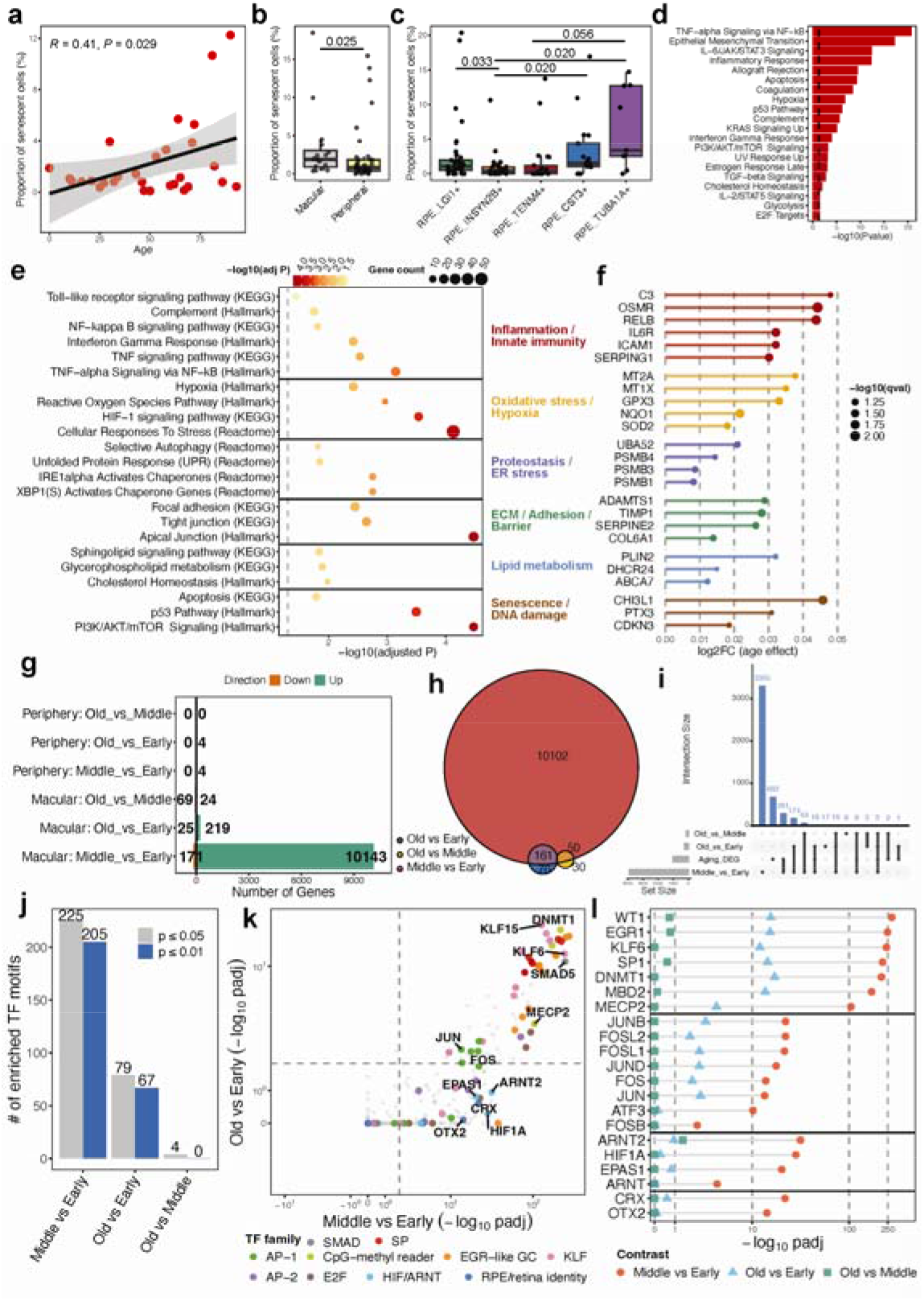
Dynamics of Cellular Senescence, Transcriptome, and Chromatin Accessibility in the Aging RPE. **a,** Scatter plot illustrating the positive correlation between age and the proportion of senescent RPE cells. **b,** Boxplot comparing the ratio of senescent RPE cells between the macula and periphery. **c,** Boxplot comparing the senescent RPE cell ratio across defined RPE types/states. Significant differences are annotated; comparisons without indicated p-values are not significant. **d,** Pathway enrichment analysis of senescence-associated genes using MSigDB hallmark gene sets. **e,** Bubble plot of RPE-relevant pathway enrichment among age-upregulated genes, categorized into six biological themes. Dot size reflects the gene count; color indicates -log_10_FDR. Pathways were selected from KEGG, MSigDB Hallmark, and Reactome databases. The dashed line marks FDR = 0.05. **f,** Lollipop plot of representative genes exhibiting increased expression with age, organized by functional category. Log_2_FC values represent the estimated age effect from the linear mixed model. **g,** Bidirectional bar plot showing the number of age-related DARs identified across different life-stage comparisons. **h,** Venn diagram showing the overlap of DARs among three comparisons in macular RPE cells. **I,** UpSet plot illustrating the intersection between age-related DAR-associated genes and aging DEGs. **j,** Number of TF motifs enriched in differentially accessible regions for each age comparison using FDR cutoffs of 0.01 and 0.05. **k,** Scatter plot comparing motif enrichment magnitudes (significance) in Middle vs. Early and Old vs. Early comparisons. Each point represents a TF, with colors indicating distinct TF families. **l,** Lollipop plot showing the enrichment significance of selected TF modules across all three age contrasts in macular RPE cells.

Next, we investigated age-related transcriptomic shifts across all RPE nuclei and within specific subtypes, identifying 1,005 pan-RPE age-associated DEGs (aging-DEGs) (642 upregulated, 363 downregulated; Figure S11a, Table S18). Notably, aging-DEGs were most abundant in LGI1+ RPE cells (155 genes) but were sparse or absent in other subtypes (Figure S11a). Functional integration of the KEGG, MSigDB, and Reactome databases revealed that the aging RPE transcriptome shifts toward a stressed, inflammatory, and metabolically remodeled state. This “aging signature” is characterized by enhanced innate immune/complement activity, oxidative stress defense, proteostasis/autophagy demand, and ECM remodeling (Figure 5e). Among the upregulated aging-DEGs, 20 are known AMD risk factors, including *C3*, *IL6R*, *OSMR*, *SOD2*, *ICAM1*, *MFRP*, *CDHR1*, and *SERPING1* (Figure 5f, Figure S11b).

Finally, we assessed age-associated chromatin accessibility changes across four life stages (pediatric: <20; early adulthood: 20–40; middle adulthood: 40–60; and old: >60 years). This analysis revealed only 0–4 DARs in the peripheral RPE but between 93 and 10,314 DARs in the macular RPE across comparisons (Figure 5g, Figure S11c). Strikingly, the early-to-middle adulthood transition exhibited much larger number of DARs, while the middle-to-old transition yielded only 93 DARs (Figure 5g), with 211 DARs shared across at least two comparisons (Figure 5h). We mapped age-related DARs to 3,911 genes, including AMD risk loci HTRA1, C3, and APOE, with 355 genes overlapping with the aging-DEGs (Figure 5i). Motif enrichment revealed the highest number of TF motifs in the early-to-middle adulthood transition (205 motifs, Padj ≤ 0.01, Figure 5j), which was dominated by GC-box/CpG-island factors, AP-1, and HIF components, alongside altered enrichment for RPE-identity TFs such as OTX2 and CRX (Figures 5k, l).

### Spatial and Multi-omic Characterization of Choroidal Endothelial Heterogeneity and Age-Associated Regulatory Remodeling

Endothelial cells (ECs) are a widespread and essential cell class that has been extensively characterized via scRNA-seq in various tissues, including the brain^34^, lungs^35^, and the uveal tract^18^. Within the choroid, we identified five vascular EC types—each showing high sequencing quality (Figure S12a) and consistent multi-donor contributions—corresponding to the vascular zonation axis: arterial, arteriole, fenestrated capillary, venule, and vein (Figure 6a). The cell proportions of these five EC types were highly consistent between the snRNA and snATAC datasets, with fenestrated capillaries constituting the highest proportion in both modalities (Figure 6a). These EC types were uniformly distributed across the macular and peripheral regions (Figure 6b) and exhibited a continuous transcriptional transition along the arterio-venous axis (Figure 6a). Notably, our snATAC-seq data successfully captured all five EC types, demonstrating strong concordance between marker gene promoter accessibility and gene expression (Figures 6a, c). We identified 1,732 marker peaks (FDR ≤ 0.05, logFC ≥ 0.5) that displayed a transitional epigenetic profile across the zonation axis (Figure S12b). This gradual shift in peak accessibility from arterial to venous systems provides compelling epigenetic evidence for vascular zonation in the human choroid. Finally, we mapped three major vascular categories within our Xenium spatial transcriptomics data: arterial/arteriole, fenestrated capillary, and venule/vein (Figure S12c). Spatial analysis confirmed that fenestrated capillaries maintained the closest proximity to the RPE across both central and peripheral regions (Figure 6d).

**Figure 6.**
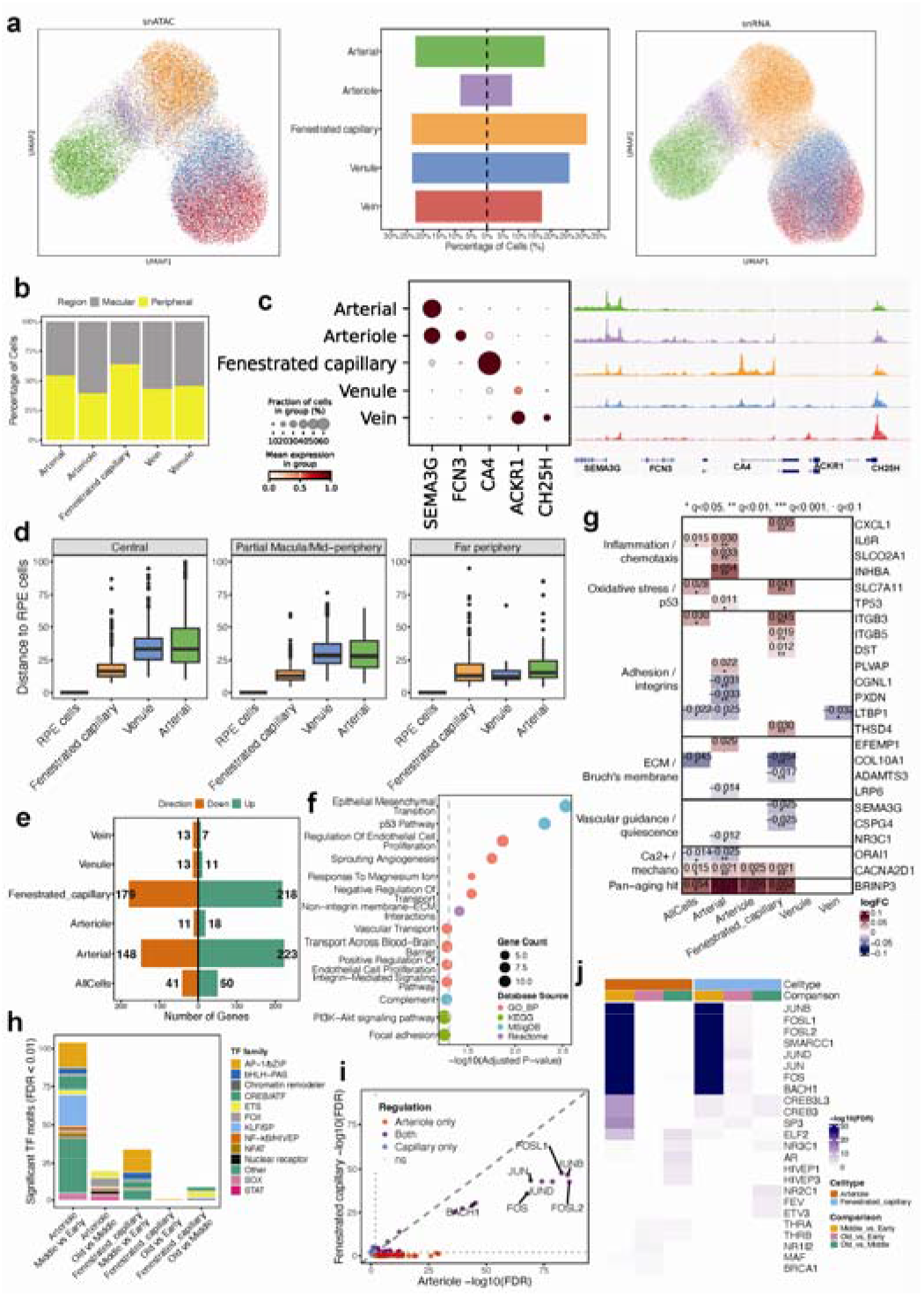
Spatial and Multi-omic Characterization of Choroidal Endothelial Heterogeneity and Age-Associated Regulatory Remodeling. **a,** UMAP visualization of choroidal endothelial subtypes based on snATAC-seq (left) and snRNA-seq (right) modalities. Middle: Bidirectional bar plot showing the relative proportion of each endothelial subtype between the two modalities. **b,** Stacked bar plot illustrating the relative distribution of cells from macular versus peripheral regions within each endothelial type. **c,** Integrated visualization of marker gene expression and chromatin accessibility. Left: Dot plot showing the mean expression and fraction of cells expressing canonical markers across endothelial subtypes. Right: Pseudobulk accessibility tracks (snATAC-seq) centered at the promoter regions of the corresponding marker genes. **d,** Boxplots showing the spatial distribution of distances between choroidal endothelial subtypes and the RPE layer, derived from Xenium spatial data. Results are faceted by anatomical region: central, partial macula/mid-periphery, and far periphery. **e,** Bidirectional bar plot displaying the number of significantly upregulated (green) and downregulated (orange) age-associated differentially expressed genes (DEGs) across all endothelial cells and specific subtypes. **f,** Dot plot of biological processes and signaling pathways enriched among aging-associated DEGs upregulated in fenestrated capillaries. **g,** Heatmap showing the effect size (logFC) of curated aging-associated DEGs. Only DEGs meeting a significance threshold of q < 0.1 are colored. **h,** Stacked bar plot providing an overview of age-associated transcription factor (TF) motif enrichment (FDR < 0.01) within choroidal endothelial subtypes. **i,** Scatter plot comparing TF motif enrichment (-log_10_FDR) between arterioles and fenestrated capillaries for the “middle adulthood vs. early adulthood” age contrast. **j,** Heatmap of top-ranked TF motifs enriched across endothelial cell types and various age-group comparisons.

We compared gene expression profiles between the macula and periphery across all EC subtypes. Regional comparisons revealed that venules and fenestrated capillaries exhibited the greatest transcriptomic differences (with 183 and 138 DEGs, respectively), whereas only 0–18 DEGs were identified in arterial, arteriole, and vein ECs (Figure S12d, Table S19). Specifically, macular venule ECs upregulated genes enriched for blood vessel morphogenesis and EC differentiation (Figure S12e), while macular fenestrated capillaries showed significant enrichment only in positive regulation of calcium ion import and regulation of leukocyte tethering or rolling. Despite these transcriptomic shifts, regional DAR analysis yielded only 2 DARs for fenestrated capillaries and 33 for arterioles, with no DARs identified in arterial, venule, or vein ECs (Table S20).

We next conducted cell composition analysis across all EC subtypes, revealing a significant negative correlation between macular venule EC proportion and age (R = -0.47, P = 0.021). Meanwhile, the peripheral vein EC ratio showed a non-significant positive trend (R = 0.35, P = 0.11; Figure S13a). To characterize transcriptomic shifts during aging, we identified 397 aging-DEGs in fenestrated capillaries (218 up, 179 down) and 371 in arterial ECs (223 up, 148 down), with substantially fewer aging-DEGs in other subtypes (Figure 6e, Table S21). Among these, 112 aging-DEGs were dynamically regulated across two or more subtypes (Figures S13b,c). Notably, *RANBP17* was progressively downregulated along age across all vascular analyses (Figure S13c). Conversely, several genes displayed age-related upregulation, most prominently *BRINP3*, which increased in arterial, arteriole, and fenestrated capillary subsets (Figure S13c). Fenestrated capillary aging-DEGs were enriched for inflammation, angiogenesis, EC proliferation, EMT, and structural remodeling, whereas arterial DEGs showed no significant pathway enrichment. Interestingly, our aging-DEG sets included genes linked to AMD pathophysiology, such as *CXCL1*, *EFEMP1*, and *LTBP1* (Figure 6g).

Lastly, we explored age-related changes in chromatin accessibility. Age-associated DAR analysis identified between 0 and 709 positively correlated DARs across EC subtypes (Table S22); the highest number of aging-DARs was observed in arteriole ECs, followed by fenestrated capillaries (Figure S13d). Motif enrichment analysis concentrated primarily on the early-to-middle adulthood transition. In arterioles, 104 significant motifs were observed in the middle-vs-early comparison, compared to only 20 in the old-vs-middle comparison. Fenestrated capillaries showed a similar trend (34 vs. 9 motifs, respectively). Motif enrichment in both subtypes was dominated by AP-1/bZIP-related factors during the early-to-middle adulthood transition (Figures 6h-j). In arterioles, the strongest motifs included FOSL2, JUNB, FOSL1, SMARCC1, JUND, JUN, and FOS. Fenestrated capillaries displayed a matching core pattern, albeit with lower magnitude, with 24 significant motifs shared between the two cell types. Collectively, our results suggest that the primary regulatory transition in the choroidal vasculature occurs between early and middle adulthood, followed by a qualitatively distinct and smaller transition from middle to old age (Figure 6j).

### Heterogeneity and Molecular Characterization of Choroid Stromal Cells

Within our snRNA-seq atlas, we profiled 185,086 fibroblasts and 24,102 mural cells. Characterization of these stromal populations identified 11 fibroblast types/states, two pericyte populations (*ADAMTS2*+, *CPM*+), and two smooth muscle cell types (vascular and *DGKI*+ ciliary) (Figures 7a-b). All defined stromal clusters exhibited high-quality metrics (Figure S14a), consistent contributions from multiple donors (Figure S14b), and robust type-specific marker gene expression (Figure 7b). Notably, five fibroblast clusters showed biased regional distributions along the macular-peripheral axis (Figure S14c). To explore their functional roles, we identified marker genes, yielding between 400 and 2,346 cluster-specific genes per population (FDR ≤ 0.05, |log_2FC| ≥ 1; Figure 7c, Table S23). Pairwise Jaccard similarity analysis indicated that most clusters are transcriptionally distinct (Jaccard index < 0.15), although C3+ and IL6+ fibroblasts emerged as closely related states (Figure S14d). GO enrichment analysis highlighted shared biological processes, such as ECM organization in fibroblasts and muscle contraction in mural cells (Figure 7d). We also identified state-specific functions: *IL6*+ and *C3*+ fibroblasts were enriched for inflammatory response and cell migration, while *FGF7*+ and *SOX6*+ subtypes were associated with nervous system development (Figure 7d).

**Figure 7.**
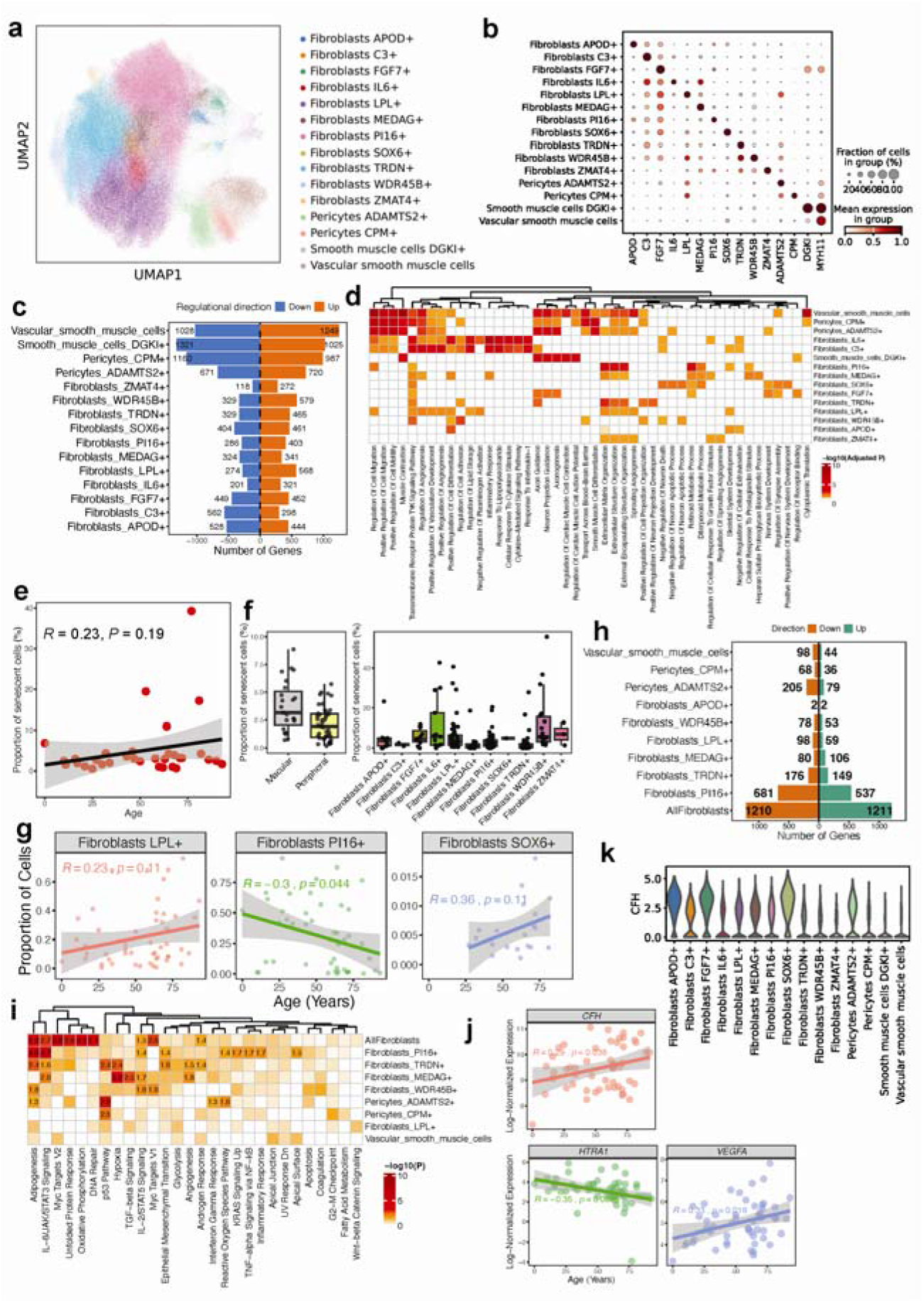
Heterogeneity and Molecular Characterization of Choroidal Stromal Cells. **a,** UMAP visualization of fibroblast and mural cell subtypes based on snRNA-seq data. **b,** Dot plot illustrating the expression profiles of marker genes across stromal subtypes. **c,** Bidirectional bar plot showing the number of significantly upregulated (orange) and downregulated (blue) subtype-specific differentially expressed genes (DEGs) identified for each population. **d,** Heatmap showing the top five enriched biological processes for each stromal subtype. **e,** Scatter plot illustrating the positive correlation between donor age and the proportion of senescent fibroblast cells. **f,** Boxplots comparing the proportion of senescent cells between macular and peripheral regions (left), and across the defined stromal subtypes and states (right). **g,** Scatter plots showing the correlation between donor age and the relative cell proportions of *LPL*+, *PI16*+, and *SOX6*+ fibroblast subtypes. **h,** Bidirectional bar plot showing the number of age-related DEGs (upregulated in green; downregulated in orange) identified across different stromal populations. **i,** Heatmap showing the top enriched biological pathways associated with aging-related DEGs for selected stromal subtype. **j,** Scatter plots showing the correlation between donor age and log-normalized expression of AMD-associated genes *CFH*, *HTRA1*, and *VEGFA* in fibroblasts. **k,** Violin plot illustrating the heterogeneous expression levels of *CFH* across the various stromal subtypes and states.

We next investigated fibroblast senescence, observing a positive, albeit non-significant, correlation between the proportion of senescent fibroblasts and donor age (R = 0.23, P = 0.19) and trended higher in the macula (P = 0.093, Figure 7e). The senescent cell ratio varied by subtype: *WDR45B*+, *IL6*+, and *FGF7*+ fibroblasts showed higher proportions, while *C3*+, *TRDN*+, and *MEDAG*+ subtypes exhibited lower rates (Figure 7f). Additionally, donor age correlated positively with *LPL*+ and *SOX6*+ fibroblast proportions and negatively with the *PI16*+ population (R = -0.3, P = 0.044; Figure 7g).

Finally, we analyzed aging-DEGs (q value ≤ 0.1) across stromal populations. The “pan-fibroblast” group exhibited the most extensive shifts, with 1,211 upregulated (enriched for RNA processing, ribosome biogenesis, and oxidative phosphorylation) and 1,210 downregulated aging-DEGs (Figures 7h-I, Table S24). Among specialized subtypes, *PI16*+ fibroblasts underwent the most substantial remodeling (537 up, 681 down), while *MEDAG*+ and *TRDN*+ subsets showed more modest profiles. In contrast, mural cells displayed fewer DEGs, with *CPM*+ pericytes showing only 36 upregulated genes (Figure 7h). Individual subsets followed unique aging trajectories: aging *PI16*+ fibroblasts specifically upregulated pathways related to inflammation and lipid metabolism, such as IL-6/JAK/STAT3 signaling, adipogenesis, and diterpenoid metabolism (Figure 7i, Figure S15). Conversely, upregulated aging-DEGs in *MEDAG*+ fibroblasts was largely dedicated to macromolecule and peptide synthesis (Figure 7i, Figure S15). Interestingly, several AMD risk genes exhibited age-dependent expression changes within the pan-fibroblast population (Figure 7j). *CFH* and *VEGFA* showed age-related upregulation (R = 0.23 and R = 0.33, respectively), whereas *HTRA1* expression trended downward with age (R = -0.15). *CFH*, in particular, demonstrated significant expression heterogeneity across different fibroblast and mural cell types (Figure 7k).

### Leveraging RPE/Choroid Multi-omics for the Functional Prioritization of AMD Risk Variants

Leveraging our multi-omics atlas of the human RPE/choroid, we integrated transcriptomic and chromatin accessibility data to prioritize AMD risk variants, identify their putative target genes, and pinpoint the specific cell types in which they function. Integration of GWAS summary statistics with snRNA and snATAC profiles mapped trait enrichments to major cell types with good cross-modal concordance; for example, intraocular pressure loci were enriched in fibroblasts and endothelial cells, while outer segment thickness loci were enriched in RPE cells (Figure 8a, Figure S16a). Notably, while initial AMD loci enrichment was observed in broad RPE open chromatin regions (OCRs) rather than gene expression, high-resolution subclass analysis revealed a significant association between AMD risk loci and the *LGI1*+ and *INSYN2B*+ RPE subpopulations (Figures 8a-b).

**Figure 8.**
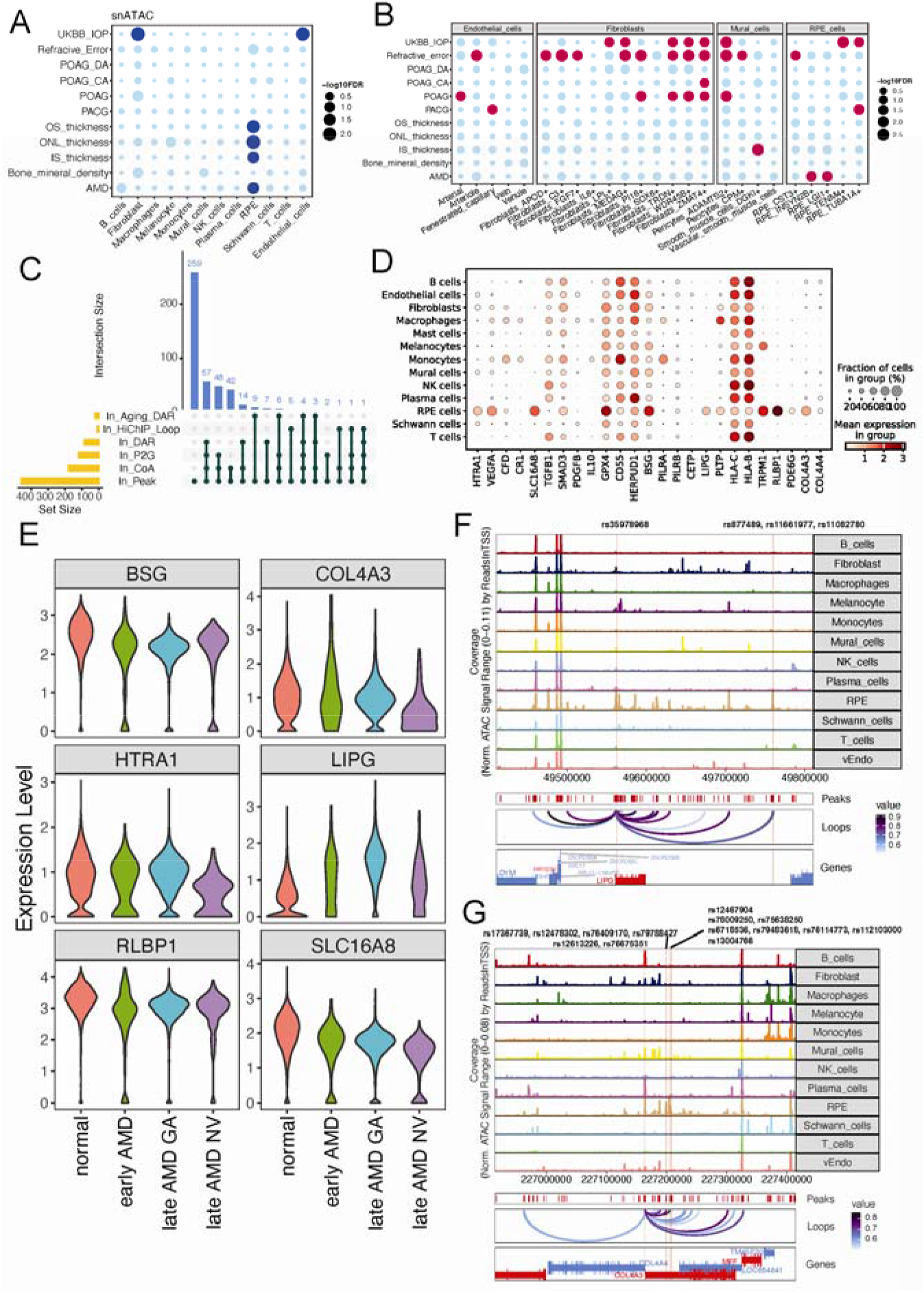
Multi-omic Integration and Functional Prioritization of AMD Risk Variants. **a,** Cell-type enrichment analysis of GWAS loci based on DARs across major cell classes. **b,** Cell-type enrichment analysis of GWAS loci based on gene expression across sub-classes. **c,** UpSet plot depicting the categorization and prioritization of AMD GWAS variants based on integrative multi-omic criteria. Peak: variants located within OCRs of RPE or choroidal cell types; CoA: variants in peaks co-accessible with other OCRs; P2G: variants in peaks significantly correlated with gene expression; DAR: variants in cell-type-specific OCRs; HiChIP loop: variants within H3K27Ac-mediated chromatin interactions; Aging DAR: variants in OCRs showing age-dependent accessibility shifts. **d,** Dot plot illustrating the expression patterns of prioritized AMD candidate genes across major cell types identified via scRNA-seq. **e,** Violin plots showing the expression levels of selected high-priority genes across RPE cells from normal donors and patients at different stages of AMD (early, late GA, and late NV). **f,** Genome track visualization of the *LIPG* locus. Highlighted regions (firebrick) indicate OCRs harboring AMD GWAS variants that are linked to *LIPG*. **g,** Genome track visualization of the *COL4A3* locus. Highlighted regions (firebrick) indicate OCRs and their associated AMD GWAS variants correlated with *COL4A3* expression.

To prioritize candidate AMD variants, we integrated 2,243 risk variants from the 95% credible sets of two landmark GWAS studies^36,37^ (Figure S16b). Of these, 459 variants (20.46%) localized within RPE/choroidal OCRs, 177 (38.56%) were co-accessible with at least one distal peak, and 119 (25.93%) resided in OCRs significantly correlated with gene expression (correlation > 0.5) (Figure 8c, Table S25). Furthermore, 86 variants were situated in cell-type-specific OCRs and 22 in age-related OCRs. This pipeline identified 120 genes putatively regulated by AMD-associated OCRs, from which we prioritized 25 candidates with existing literature support. Among these, *SLC16A8*, *LIPG*, *RLBP1*, and *COL4A3* exhibited RPE-specific expression, while *HTRA1* and *VEGFA* were expressed broadly but peaked in the RPE. Conversely, *CFD* and *PILRA* were specifically localized to monocytes and macrophages (Figure 8d, Figure S16c).

Our integrative approach both replicated and expanded upon existing functional annotations. At the *ARMS2/HTRA1* locus, where previous studies identified a single risk variant in an RPE-specific OCR^38^, we identified six variants across two RPE OCRs correlated with *HTRA1* expression (Figure S16d). Leveraging the inclusion of AMD donor cells across various disease stages in our integrated scRNA-seq atlas, we evaluated the expression of candidate genes from early through late-stage AMD. We observed a significant downregulation of *HTRA1* in the RPE of neovascular AMD (nAMD) donors compared to normal controls (Figure 8e). Similarly, for *VEGFA*, we identified four additional OCRs harboring nine GWAS variants correlated with its expression, including rs943080 previously reported by Wang et al.^24^ (Figure S16e). We also identified high-priority risk genes with novel regulatory links. For *LIPG*, we identified four GWAS variants across two OCRs (one of which is RPE-specific) correlated with its expression (Figure 8f). Notably, *LIPG* exhibited significant upregulation in RPE cells across all stages of AMD (Figure 8e). For *COL4A3*, an essential component of Bruch’s membrane, we identified six OCRs (two RPE-specific) harboring 14 GWAS variants correlated with its expression (Figure 8g). *COL4A3* expression was significantly decreased in RPE cells from AMD patients, with the lowest levels observed in late-stage nAMD (Figure 8e). We similarly identified variant-harboring OCRs for *BSG*, *RLBP1*, and *SLC16A8*, all of which were downregulated in RPE cells from AMD donors (Table S25).

## Discussion

In this study, we assembled a comprehensive multi-modal single-cell atlas of the human RPE/choroid, integrating transcriptomic data from 719,813 cells and nuclei (scRNA-seq/snRNA-seq) and epigenomic data from 234,007 nuclei (snATAC-seq), alongside seven Xenium spatial slides. This dataset, derived from 102 multi-ancestry donors across a broad age range, captures cellular composition, transcriptomic states, chromatin accessibility, and spatial positioning across the macular and peripheral regions throughout the adult lifespan. Our integrated atlas confirms the major cell classes of the RPE/choroid complex while significantly increasing resolution. Sub-clustering of the snRNA-seq compartment yielded 48 cellular subtypes and states; notably, the recovery of 68,871 RPE nuclei provides approximately an order of magnitude more RPE cells than prior public scRNA-seq studies combined^13,15,16,18,19,22,23^. We also report a systematic comparison between scRNA-seq and snRNA-seq modalities. While major cell-class identities are well-aligned across platforms, the detection of an average of 6,438 differential detection genes per class indicates that direct cross-modality interpretation of expression magnitude requires caution. This resource provides a high-resolution reference for ocular homeostasis and a framework for prioritizing causal genes and cell types underlying AMD.

Compared with prior RPE/choroid transcriptomic studies, our atlas recapitulates all previously described cell types and resolves a substantially greater degree of cellular heterogeneity, including five RPE types/states, eleven fibroblast subclasses, and two distinct pericyte populations. The RPE has traditionally been treated as a homogeneous monolayer or stratified by macular versus peripheral location. Recent morphometric work^39^ proposed five structural RPE classes (P1–P5), but their transcriptomic correlates remained undefined. Here, we identify five RPE clusters that segregate along a central-to-peripheral axis and are marked by *LGI1*, *INSYN2B*, and *TENM4*. Two additional clusters (*CST3*⁺ and *TUBA1A*⁺) display stress-response signatures and elevated senescence markers, suggesting they represent specialized reactive states rather than stable subtypes.

We further show that fibroblast subclasses exhibit distinct regional distributions, age-associated compositional shifts, and variable *CFH* expression. Given the anatomical proximity of fibroblasts to the fenestrated choriocapillaris and adjacent choroidal cell types, we hypothesize that CFH^hi^ fibroblasts establish a local protective niche that restrains complement activation and chronic inflammation. Within the LPL+ fibroblast population, we also observed a subset of cells expressing *LGR5*. These *LGR5*+ cells are established coordinators of replication, differentiation, and tissue homeostasis across multiple organs^40–42^. Dysfunction of *LGR5*+ stromal cells is strongly linked to scleroderma^43^, a progressive multi-organ fibrotic disease. Future spatial mapping of the RPE-choroid complex will define the topography of these *LGR5*+ fibroblasts and their potential relevance to macular pathology. The specific expression of *VEGFA* in *C3*⁺ fibroblasts further implicates this subset in the active modulation of choroidal angiogenesis. Together, these findings argue that choroidal fibroblasts function as active regulators of local tissue homeostasis rather than passive structural scaffolds. Spatial validation will be essential to define the precise co-localization patterns and multicellular communication networks that govern these newly resolved subtypes. In summary, this study provides a healthy baseline reference for disease comparison, grounded in the premise that ocular tissues are not biologically uniform and that disease does not affect all cells equally. By reframing the posterior eye from a static tissue to a dynamic cellular ecosystem, this atlas offers a foundational blueprint for future high-resolution spatial studies.

As the primary risk factor for AMD^44^, aging induces numerous phenotypic changes in healthy RPE/choroid tissues. these include decreased choriocapillaris density, increased flow deficits, reduced choroidal thickness, increased Bruch’s membrane thickness, and elevated RPE autofluorescence^45–48^. However, the underlying molecular and cellular mechanisms remain largely unknown. Furthermore, our understanding of cellular compositional changes in the RPE across the lifespan has been limited by contradictory results^49^. While research confirms that RPE density is region-dependent—peaking in the fovea and decreasing toward the periphery^48,50,51^—reports on age-related density changes vary. Here, we present the age-associated compositional, transcriptomic, and epigenetic changes in the RPE/choroid, providing potential molecular and cellular mechanisms for the clinical and morphometric observations.

We demonstrate that while central and mid-peripheral RPE populations exhibit a modest decrease with age, the far-peripheral *TENM4*+ RPE population significantly increases. We also observed age-associated shifts in the proportions of both immune and non-immune cells. For instance, the decline in melanocytes aligns with reported age-related decreases in skin melanocyte density, estimated at 10–20% per decade after age 30^52^. Within the stroma, the age-related cell loss is concentrated in *PI16*+ fibroblasts—one of the largest fibroblast populations (30.75%)—and macular venule ECs. Collectively, these findings suggest that the clinical observation of reduced choroidal thickness with age may be driven by the coordinated loss of melanocytes, venule ECs, and *PI16*+ fibroblasts.

At the transcriptomic level, aging induces common inflammatory shifts across multiple cell types, yet with distinct cell-type specificity. In the stroma, remodeling is concentrated in *PI16*+ fibroblasts (537 upregulated, 681 downregulated genes), which exhibit inflammatory and lipid-metabolic signatures, including the IL-6/JAK/STAT3 and adipogenesis pathways. Notably, several AMD risk genes (*CFH*, *VEGFA*, *HTRA1*) show age-dependent expression within the pan-fibroblast pool. Simultaneously, the RPE shifts toward complement and innate immune activation, oxidative stress defense, proteostasis demand, and ECM remodeling, with 20 known AMD risk genes upregulated. Together with the regional accumulation of senescent RPE cells in the macula, these signatures support a model in which AMD susceptibility reflects accumulated regulatory and senescence-associated changes initiated in early adulthood and concentrated at the macula. The significant age-associated enrichment of genes related to ER stress, proteostasis, and lipid metabolism within the RPE likely represents a compensatory that ultimately becomes insufficient to counteract the cumulative cellular stress. The upregulation of lipid processing pathways likely reflects the metabolic burden imposed by the continuous clearance of outer segment lipids and toxic retinoid byproducts, while the concurrent induction of ER stress markers underscores the proteostatic strain caused by non-degradable protein aggregates. These molecular signatures provide a transcriptomic basis for the elevated autofluorescence observed in aged RPE, marking a transition toward cellular senescence and metabolic exhaustion.

Fenestrated capillaries exhibit the most pronounced age-associated transcriptomic divergence among ECs (397 DEGs), enriched for inflammation, angiogenesis, EMT, and structural remodeling. Crucially, this shift was characterized not only by a pro-angiogenic signature but also by a significant enrichment in the negative regulation of transport. This suggests that while angiogenesis-related turnover may maintain overall endothelial cell density—consistent with our observation that the fenestrated EC ratio remains stable with age—the resulting vascular bed is likely functionally compromised. The downregulation of transport mechanisms, paired with the upregulation of structural remodeling and EMT genes, points to a loss of the specialized exchange capacity required to support RPE homeostasis. We propose that this ‘molecular sealing’ of the aging choriocapillaris contributes to clinically observed flow deficits^46^, representing a state of attenuated vascular competence without overt loss of vascular presence. This is consistent with the choriocapillaris dropout reported in AMD histopathology^53^.

Finally, we explored the epigenetic aging signal, which proved to be highly tissue-compartment specific. In the RPE, ECs, and fibroblasts, we observed thousands of aging-DARs in the macula, compared to a maximum of only a few DARs in the periphery. In the macular RPE, the most substantial chromatin remodeling occurs between early and middle adulthood (up to 10,314 DARs), with comparatively little remodeling occurring during the transition from middle-to-old age (93 novel DARs). A similar pattern was observed in ECs: motif enrichment is concentrated in the early-to-middle window in arterioles and fenestrated capillaries, with AP-1/bZIP as the dominant signal. This early-adulthood reorganization precedes the typical clinical onset of AMD by decades, pointing to a critical regulatory window in which the molecular substrate of later disease is established.

Integration with AMD GWAS data underscores the practical utility of this resource. MAGMA (transcriptomic) and LDSC (chromatin) enrichments pinpoint the *LGI1*+ and *INSYN2B*+ RPE subpopulations as having significantly higher genetic risk for AMD. Our pipeline links risk variants to 120 putative target genes, replicating known loci (*HTRA1*^38^, *VEGFA*^24^) at higher resolution while expanding the set of supported targets to include *LIPG* and *COL4A3*. *COL4A3* encodes the α3chain of the specialized α3α4α5(IV) collagen IV network. This network is biologically distinct from the ubiquitous α1α1α2(IV) network and is critical for the structural integrity of basement membranes at filtration and stress-bearing surfaces, including the ocular basement membranes affected in Alport syndrome^54,55^. The downregulation of *COL4A3* reflects a failure to synthesize this specialized, anti-angiogenic basement membrane program. Conversely, Bruch’s membrane (BrM) thickening represents the failed clearance of inert and pathogenic debris; these two phenomena describe distinct, complementary facets of ECM dyshomeostasis that are expected to co-occur in the aging RPE^56^. In neovascular AMD the combination of a thickened, poorly permeable BrM and reduced α3(IV)-mediated maintenance likely weakens a critical endogenous brake on choroidal endothelial invasion.

*LIPG* encodes endothelial lipase, a member of the triglyceride lipase family with high phospholipase activity toward HDL phospholipids^57,58^. Genetic and human-sequencing studies show that reduced *LIPG* activity raises circulating HDL-C, establishing *LIPG* as a bona fide regulator of HDL metabolism^59–61^. Large GWAS and follow-up studies consistently implicated HDL-associated loci, including *LIPC*, *CETP*, and *ABCA1*, in AMD susceptibility^36,62–65^. In parallel, pathology studies showed that BrM and drusen accumulate cholesterol-rich, apoB- and apolipoprotein-containing particles with age, supporting the hypothesis that defective lipid handling at the RPE–BrM interface contributes directly to lesion formation^66–71^. In this context, the upregulation of *LIPG* in the RPE is attractive not because it recapitulates plasma lipid physiology, but because it could reshape local lipoprotein and phospholipid homeostasis in a tissue compartment already prone to age-dependent lipid retention. Together, these genes, alongside prioritized cell-type contexts, represent concrete starting points for future functional follow-up.

While this study provides a comprehensive map of RPE/choroidal organization, it also establishes a roadmap for subsequent high-resolution investigations. Based on these findings, future efforts utilizing high-resolution spatial multi-omics or whole-mount imaging will be essential to map the spatial architecture and cellular neighborhoods of these populations. Although our initial Xenium panel was optimized for the neural retina, this atlas now provides the precise marker genes required to design next-generation spatial probes tailored specifically to the RPE and choroid. Such targeted panels will be instrumental in resolving the spatial niches of closely related populations, such as the various fibroblast subclasses identified here, thereby elucidating the intricate cell-cell interactions governing the RPE/choroid complex. Furthermore, while our current spatial data offers a high-resolution snapshot of an aged donor, it serves as a robust baseline for future studies spanning a broader age range to further enhance generalizability. Finally, although the number of AMD-derived RPE cells in current datasets is limited, the integrated framework of this atlas provides a powerful foundation for the targeted inclusion of additional disease samples. By expanding this resource, future work can leverage our high-resolution annotations and variant-to-gene mappings to dig deeper into the specific molecular mechanisms driving AMD. Together, this atlas is a concrete starting point for cell-type-and region-specific therapeutic strategies in ocular aging and disease.

## Methods

### Ethics, Sample Collection, and Tissue Dissection

This study was conducted in accordance with all relevant ethical regulations and the tenets of the Declaration of Helsinki. The study protocol was reviewed and approved by the Institutional Review Boards (IRBs) at the University of California, Irvine; Baylor College of Medicine; the University of Utah; and the State University of New York at Buffalo. Human donor eyes were obtained post-mortem from the Baylor Lions Eye Bank, the Utah Lions Eye Bank, the Lions World Vision Institute, and the Lions Eye Bank of Texas. Informed consent was obtained from all donors or their legal representatives, and all samples were de-identified in compliance with HIPAA Privacy Rules prior to receipt.

RPE/choroid samples were collected within 10 hours post-mortem for transcriptomic profiling and within 19 hours post-mortem for snATAC-seq profiling. Donor medical histories were thoroughly reviewed to ensure that only individuals without evidence of retinal pathology were included in the study. Using 2 mm and 6 mm disposable biopsy punches, the foveal and macular regions were dissected. The 2 mm foveal centralis was excluded, while the surrounding 6 mm macular ring (macula) was retained for profiling. Peripheral RPE/choroid tissues were subsequently collected from the posterior segment following a ‘butterfly’ incision. For snRNA-seq, all collected tissues were immediately flash-frozen in liquid nitrogen and stored at -80°C until nuclei extraction and downstream processing.

### Single-cell/nucleus preparation, library construction, and sequencing

For scRNA-seq, isolated RPE/choroid sheets were incubated in pre-warmed Trypsin-EDTA/DNase I digestion buffer and enzymatically digested at 37°C for 20-45 min, with intermittent gentle pipetting until a single-cell suspension was achieved. The suspension was then filtered through a 50-70 µm cell strainer and neutralized using Hibernate A buffer supplemented with 20% FBS. Cells were collected via centrifugation at 300x g for 5min, the supernatant was removed, and the cell pellet was washed once with cold Hibernate A buffer. The final pellet was resuspended in 200 ul of cold Hibernate A buffer.

Nuclei for snRNA-seq were isolated using the GentleMacs Octo Dissociator with C-tubes in ice-cold Nuclei Extraction Buffer. The resulting suspension was filtered through 70-µm MACS SmartStrainers and centrifuged at 300 × g for 5 minutes at 4°C. The nuclei pellet was resuspended in wash buffer (10 mM Tris–HCl, 10 mM NaCl, 3 mM MgCl₂, 1% BSA), gently triturated to dissociate residual tissue fragments, and passed through 30-µm pre-separation filters into 5-ml round-bottom tubes. A second centrifugation at 500 × g for 5 minutes at 4°C, the nuclei were resuspended in PBS for downstream processing.

Single-cell and single-nucleus gene expression libraries were generated using the Chromium Next GEM Single Cell 3’ v3.1 Reagent Kit (10x Genomics). Single-nucleus ATAC libraries were prepared following the Chromium Next GEM Single Cell ATAC v1.1 protocol. All sequencing was performed on the Illumina NovaSeq 6000 platform.

### Data Preprocessing and Quality Control

Raw sequencing reads from both curated public datasets and our newly generated data were processed using Cell Ranger (v7.1.0) and aligned to the 10x Genomics Human GRCh38-2020-A reference genome. To ensure technical consistency, we employed the cellqc pipeline^11^ (https://github.com/lijinbio/cellqc), an integrated workflow that automates ambient RNA correction, doublet detection, and cell filtering. Within the cellqc framework, ambient RNA contamination was corrected using SoupX^68^, and potential doublets were systematically identified and removed via DoubletFinder^69^. Following these automated corrections, we applied standardized hard-filtering criteria to the resulting feature-barcode matrices. Specifically, we retained only those cells and nuclei that met the following quality thresholds: a minimum of 500 total UMIs, at least 300 unique genes, and a mitochondrial transcript fraction below 10% for scRNA-seq or 5% for snRNA-seq. This unified approach ensured a high-quality, standardized dataset for subsequent atlas construction.

### Xenium data generation

Ocular tissues were collected from a 77-year-old female European donor with no recorded history of ophthalmic disease and a post-mortem interval (PMI) of 3.2 hours. To preserve tissue morphology and RNA integrity, tissues were fixed for 48 hours. The sample was fixed in 4% paraformaldehyde (PFA, Catalog# 50980487) supplemented with 5% sucrose, while the remaining samples were fixed in Modified Davidson’s Fixative (Catalog# 64133-50). To enhance fixative penetration, several small incisions were made at the pars plana prior to immersion fixation. Following fixation, tissues were stored short-term in either 1% PFA/5% sucrose or 70% ethanol before paraffin processing. For embedding, tissues were dehydrated through a graded ethanol series (70%, 80%, three changes of 95%, and two changes of 100% ethanol). Samples were then cleared in three changes of Histo-Clear and infiltrated with two changes of molten paraffin. Each dehydration, clearing, and infiltration step was performed for one hour. Following infiltration, tissues were trimmed to fit Xenium slides, embedded in paraffin blocks, and stored at −20°C until sectioning.

Formalin-fixed paraffin-embedded (FFPE) tissues were sectioned horizontally at 5 μm thickness using a Leica RM2255 microtome and mounted onto Xenium slides, following the 10x Genomics protocol (CG000578_RevE). Sections were deparaffinized and subjected to RNA decrosslinking (protocol CG000580). Probe hybridization was performed for 16–24 hours using a customized Xenium Human Eye Panel targeting 478 genes (Design ID: G33B7Z). Subsequent enzymatic ligation and rolling circle amplification were performed to enable the detection of spatially resolved transcripts. Following amplification, cell segmentation staining and background fluorescence quenching were carried out according to the manufacturer’s instructions (10x Genomics, CG000749 Rev B). Images were acquired and transcripts were decoded on the Xenium Analyzer. The resulting outputs included cell-by-gene expression matrices and transcript-level spatial coordinates for each tissue section.Following data acquisition, the same tissue sections were stained with hematoxylin and eosin (H&E) according to the manufacturer’s protocol (CG000613 Rev B, 10x Genomics). Post-Xenium H&E images were acquired by using Keyence BZ-X810 and were used to assess tissue morphology, support quality control, and refine image-based cell segmentation.

### Integration, clustering, and annotation

To integrate datasets across different donors and studies while minimizing batch-driven technical variation, we employed scVI^25^. Integration was conducted independently for the scRNA-seq and snRNA-seq cohorts to account for the inherent technical differences between whole-cell and single-nucleus libraries. Integration was performed first on the full dataset and subsequently for each major cell class—including RPE, endothelial, immune, mural, fibroblasts, melanocytes, and Schwann cells. Highly variable genes (HVGs) were identified using the Seurat flavor, and the top 10,000 HVGs were retained as input features for the model. The scVI model was trained to learn a shared low-dimensional latent space (n_layers=2, n_latent=30), and the resulting embeddings were utilized for graph-based clustering via the Leiden algorithm^72^. To determine optimal cluster granularity, we implemented a two-level hierarchical clustering strategy. In the first level, Leiden resolutions between 0.1 and 0.5 were evaluated to identify major transcriptional groups while preventing over-clustering. Clusters that remained transcriptionally heterogeneous were subjected to a second round of Leiden clustering (resolutions 0.1–0.4) to resolve distinct subclusters.

Major cell classes were first identified using canonical marker genes, with the immune cell population initially annotated as a single broad class based on *PTPRC* expression. Other major classes were defined by the following markers: *RPE65* and *BEST1* (RPE); *PECAM1*, *VWF*, *PTPRB*, and *FLT1* (Endothelial); *PDGFRA* and *FBLN1* (Fibroblasts); *MLANA*, *TYR*, *PAX3*, and *TEX41* (Melanocytes); *PDGFRB* and *ACTA2* (Mural cells); and *NRXN1* and *MBP* (Schwann cells). Following this initial classification, the broad immune population was further resolved into specific cell types using reference mapping with CellTypist^73^. For all other major classes, we performed independent sub-clustering and applied the same marker-based strategy to assign high-resolution cell-type labels. NS-Forest^26^ was utilized to identify de novo marker genes. Final cell-type assignments were determined based on a consensus of de novo markers, canonical gene expression (where applicable). Quality control metrics (nCounts, nGenes, and mitochondrial transcript fraction) were confirmed to be consistent across all annotated cell types; furthermore, each major cluster was supported by cells from multiple donors to ensure results were not driven by individual-specific biases.

### Xenium data processing and spatial analysis

Spatial transcriptomic profiling was performed using Xenium Ranger (v3.0; 10x Genomics) with a custom-designed panel comprising 478 genes (Table S5). Initial data processing was conducted using Squidpy^74^, and cells with fewer than 20 detected transcripts were excluded. Raw counts from all segmented cells across all slides were batch-corrected and integrated using scVI, following the workflow described above. Cell class annotation was performed using the aforementioned canonical marker genes. Cells annotated as neural retinal populations were excluded from downstream analysis. To ensure the spatial specificity of our RPE/Choroid analysis, cells from the optic nerve and optic nerve head were manually identified and removed using the Xenium Explorer 3 selection tool. The spatial distance between endothelial cells and the RPE was estimated using the sq.tl.var_by_distance() function in Squidpy.

To validate the accuracy of cell-type assignments in the Xenium spatial dataset, we performed a co-embedding analysis with the integrated sc/snRNA-seq RPE/Choroid atlas. The Xenium and sc/snRNA-seq datasets were merged and processed within the Seurat (v5)^75^. Following log-normalization, we performed modality-aware scaling (ScaleData) by regressing out variation associated with the data modality. Principal Component Analysis (PCA) was performed on the shared feature space, followed by cross-modality batch correction using Harmony^76^. A joint Shared Nearest Neighbor (SNN) graph was constructed using the first 30 Harmony dimensions, and the manifold was visualized using Uniform Manifold Approximation and Projection (UMAP).

To resolve heterogeneity within the RPE population, we implemented a marker-based classification strategy using a panel of subtype-specific genes: *EYA4*, *AC018742.1*, *UGDH*, and *CMTM6*. For each RPE cell, expression levels for these markers were extracted from the normalized Xenium count matrix. Cells were assigned to a subtype based on a highest-expression logic, where a cell was labeled according to the marker exhibiting the maximum expression value relative to the others in the panel. To maintain high specificity and mitigate the impact of background noise, an expression threshold of 0.1 was applied. Cells where the maximum expression of all markers failed to meet this threshold were categorized as “Unassigned.”

### Cross-Modality Transcriptomic Alignment and Comparative Analysis

We employed SysVI^77^ and MetaNeighbor^28^ to evaluate the transcriptional similarity and annotation consistency between the scRNA-seq and snRNA-seq modalities. To generate a shared latent space for whole-cell and single-nucleus datasets, we utilized SysVI. Integration was initiated by identifying the top 2,000 highly variable genes (HVGs) using the Seurat flavor. The SysVI model was configured with a VAMP prior (n=5) and utilized modality (scRNA vs. snRNA) as the system key, with sample ID included as a categorical covariate to regress out batch-specific technical effects. The resulting unified latent representation was used for graph-based neighbor construction and UMAP visualization to qualitatively assess concordance between modalities. To quantitatively examine cell-type similarities across technologies, raw counts were first aggregated into pseudo-bulk profiles for each major class per sample. Using this pseudo-bulk matrix, cross-dataset similarity was assessed via the MetaNeighbor R package. Specifically, HVGs were identified using the variableGenes() function, designating “dataset” (scRNA vs. snRNA) as the sample source. The mean Area Under the Receiver Operating Characteristic Curve (AUROC) was then calculated for each “majorclass” across datasets using the MetaNeighborUS() function.

To characterize specific transcriptomic differences between scRNA-seq and snRNA-seq, we performed differential detection gene (DDG) analysis for each of the 13 major cell classes using edgeR^78^. For each class, raw counts were aggregated into pseudobulk totals and filtered to remove low-expressed genes via the filterByExpr function. Libraries were normalized using the trimmed mean of M-values (TMM) method. To isolate modality-specific effects from biological and population-level noise, we utilized a generalized linear model (GLM) incorporating library type as the primary variable. The model adjusted for age, sex, and the first principal component (PC1) derived from genetic data to account for ancestry-related variations. Genes were considered significant if they met a false discovery rate (FDR) < 0.05 and a |log_2_ (fold change)| > 1. Significant genes were subsequently exported for functional enrichment analysis using Enrichr^79^.

### Annotation of snATAC-seq Data and Co-embedding with scRNA-seq

Raw snATAC-seq reads were processed using the Cell Ranger ATAC pipeline (v2.0.0) and analyzed with the ArchR^80^ package (v1.0.3). Quality control was performed by removing cells with ≤ 1,000 fragments or a transcription start site (TSS) enrichment score ≤ 4. Doublets were identified and filtered using the ArchR functions addDoubletScores() and filterDoublets() with default parameters. Dimensionality reduction was conducted using Iterative Latent Semantic Indexing (LSI), and single cells were visualized in a reduced-dimensional space via UMAP using the addUMAP() function. Major cell classes for snATAC-seq were annotated by aligning the snATAC-seq datasets with scRNA-seq data, comparing the snATAC-seq gene score matrix to the scRNA-seq gene expression matrix. Peak calling was performed using MACS2 through the ArchR addReproduciblePeakSet() function. Marker peaks for each major class were identified using the getMarkerFeatures() function with a log2 fold change ≥ 1 and FDR ≤0.05. For endothelial cell (EC) subtypes, marker peaks were defined using a more relaxed threshold of log2 fold change ≥ 0.5 and FDR ≤0.05. Co-accessible peaks were identified using the getCoAccessibility() function with a correlation cutoff of 0.3. Peak-to-gene links were established using the addPeak2GeneLinks() function by integrating scRNA-seq data, applying a correlation cutoff of 0.5, an FDR cutoff of 0.05, and a maximum distance of 250 kb.

For the multi-omic integration of sc/snRNA-seq and snATAC-seq, we employed scGLUE^81^. Cell-by-peak fragment count matrices were generated both within specific major cell classes and across the full dataset using Seurat and Signac^82^. all snATAC-seq cells were co-embedded with scRNA-seq cells in supervised mode, leveraging the pre-established major class and cell-type annotations for both modalities to ensure a high-fidelity integrated latent space.

### Identification of regulon from snATAC-seq data

To delineate enhancer-mediated regulatory programs across RPE/Choroid cell classes, we integrated our annotated snATAC-seq and snRNA-seq datasets using the SCENIC+^83^ (v1.0a2). To optimize computational performance, both modalities were downsampled to a maximum of 2,000 representative cells per major class. Within the pycisTopic environment, cis-topics were derived from the consensus peak set of the snATAC-seq data using the run_cgs_models_mallet() function. We evaluated multiple topic models via evaluate_models() to determine the optimal number of topics for our dataset. Regulatory networks centered on transcription factors (TFs)—including candidate distal regulatory elements and their associated target genes—were inferred using the precomputed human cisTarget database. To identify regulatory programs specific to individual cell types, we calculated eRegulon Specificity Scores.

### Regional DEGs and DARs

To characterize specific transcriptomic differences between macula and periphery, we performed differential detection gene (DDG) analysis for each of the 13 major cell classes using edgeR^78^. For each class, raw counts were aggregated into pseudobulk totals and filtered to remove low-expressed genes via the filterByExpr function. Libraries were normalized using the trimmed mean of M-values (TMM) method. To isolate modality-specific effects from biological and population-level noise, we utilized a generalized linear model (GLM) incorporating library type as the primary variable. Genes were considered significant if they met a false discovery rate (FDR) < 0.05 and a |log_2_ (fold change)| > 1. Significant genes were subsequently exported for functional enrichment analysis using Enrichr^79^. To identify region-associated differentially accessible regions (DARs) for each major cell class, the getMarkerFeatures() function from ArchR was applied to the consensus peak set. Peaks were defined as regional DARs if they met a threshold of |log_2_(fold change)| ≥ 0.5 and FDR ≤ 0.2.

### RNA velocity, trajectory inference, and pseudotime associate gene identification

Spliced and unspliced transcript counts were generated from Cell Ranger-aligned BAM files using velocyto^84^. Sample-level loom files were integrated with the filtered snRNA-seq atlas by harmonizing cell barcodes and retaining only overlapping cells; these files were subsequently concatenated into a unified spliced/unspliced matrix. RNA velocity was estimated using the scVelo^85^ framework, with velocity vectors projected onto the precomputed UMAP embedding. To summarize the transition structure among cell types, we employed Partition-based Graph Abstraction (PAGA)^86^, transferring the precomputed nearest-neighbor graph into a scVelo-compatible format to ensure consistency with the global embedding. Finally, genes associated with pseudotime were identified using the fit_models() function in Monocle3^87^, utilizing a negative binomial distribution (expression_family = “negbinomial”).

### Identification of Senescence Cells and Senescence-Associated Genes

The single-cell dataset was randomly partitioned into 29 batches while preserving the original cell-type proportions. Each batch was independently analyzed using DeepSAS^88^ to identify senescent cells (SnCs) and senescence-associated genes (SnGs). For each cell type, senescent cells identified by DeepSAS were mapped onto the gene–cell graph, and attention scores associated with gene-connected edges were extracted. A senescence-associated gene score (SnG score) was computed for each gene by averaging attention scores across all senescent cells within the corresponding cell type. To identify biologically relevant SnGs, differential expression analysis was performed separately for each cell type using the Wilcoxon rank-sum test implemented in Scanpy^89^, comparing senescent and non-senescent cells. Genes with log fold change ≥ 0.25 in senescent cells were retained and intersected with genes having non-zero SnG scores. Reproducibility across batches was assessed by quantifying the frequency of each gene–cell type pair across all batches.

### Analysis of Age-Associated Compositional, Transcriptomic, and Epigenetic Dynamics

To investigate shifts in cell type proportions throughout the human lifespan, we performed a regression-based composition analysis for selected major/sub cell classes. Cell type proportions were calculated at the donor level. To ensure statistical robustness, we excluded donors with fewer than 100 total cells in the target major/sub classes. Furthermore, specific cell-type-donor and region (if applicable) combinations were excluded if the raw cell count was less than 10. For each donor, the proportion of each sub-cluster was calculated as the ratio of sub-cluster-specific cell counts to the total number of cells recovered for that donor within the relevant major class. Age-associated trends were modeled using a linear regression. Pearson correlation coefficients and associated p-values were calculated to assess the strength and significance of these relationships.

To identify age-associated genes, differential expression analysis was conducted using the snRNA-seq dataset. For each cell type, raw read counts were aggregated into donor-level pseudobulks. Only donor samples containing at least 20 cells for a given cell type were included. Lowly expressed genes were filtered using a counts-per-million (CPM)-based threshold, retaining genes with a mean CPM ≤5 across all donors. Samples exhibiting a read-count correlation < 0.75 with more than 65% of other samples were identified as outliers and excluded. The filtered pseudobulk matrices were normalized using the Trimmed Mean of M-values (TMM) method via edgeR. We then employed the dream framework, a linear mixed-model approach implemented in the variancePartition^90^ package, to model gene expression changes as a function of age while adjusting for covariates. Age was modeled as a continuous variable, with sex and ancestry included as fixed-effects covariates and batch as a random intercept. Multiple testing correction was performed via the qvalue package; genes with q < 0.1 were considered significantly associated with age.

To identify age-associated differentially accessible regions (DARs), donors were categorized into four groups: pediatric, early adulthood, middle adulthood, and late adulthood (old). Using the early adulthood group as a reference, we utilized the ArchR getMarkerFeatures() function to identify DARs in middle and late adulthood for macular and peripheral cells separately. A peak was defined as an age-associated DAR if it exhibited a |log_2_(fold change)| ≥ 0.5 and FDR ≤ 0.1. These DARs were assigned to genes if they were located within a gene promoter or linked via peak-to-gene (P2G) links, defined by a correlation > 0.5 and an FDR ≤ 0.05.

### Cell type enrichment of GWAS loci and Prioritization of Risk Variants

We evaluated the enrichment of GWAS-associated traits within RPE/Choroid cell populations using both chromatin accessibility and gene expression profiles. To assess accessibility-based enrichment, we employed stratified linkage disequilibrium (LD) score regression via LDSC^91^ (v1.0.1) to determine if trait heritability was significantly enriched within cell-type-specific open chromatin regions (OCRs) derived from snATAC-seq data. HapMap3 SNPs were annotated based on their overlap with cell-type-specific OCRs, and LD scores were computed across 1-cM windows using the 1000 Genomes Project Phase 3 reference panel. Enrichment was estimated relative to the baseline LD model (1000G_Phase3_baselineLD_v2.2_ldscores).

For the transcriptomic analysis, we utilized MAGMA-Celltyping^92,93^ (v2.0.8) to identify positive correlations between cell-type-specific gene expression and gene-level genetic associations. GWAS summary statistics were standardized using MungeSumstats^94^. Gene-level association statistics were calculated using a genomic window of -35 kb to +10 kb around each gene, referenced against the 1000 Genomes European panel. The snRNA-seq expression matrices were processed using EWCE^95^ (v1.16.0) prior to linear enrichment testing. For both accessibility- and expression-based analyses, p-values were adjusted for multiple testing using the Benjamini-Hochberg method across all cell populations and traits.

To prioritize AMD GWAS risk variants, we compiled a union set of variants from 95% credible sets identified in two landmark studies^36,37^. These risk variants were intersected with the genomic coordinates of several regulatory features: all open chromatin regions (OCRs), major cell-type-specific DARs, co-accessible peaks, and age-associated DARs. Risk variants were assigned to candidate genes if they overlapped with a gene promoter or were situated within a distal peak linked to a gene via peak-to-gene (P2G) links (defined by a correlation > 0.5 and an FDR ≤ 0.05). Genomic track visualizations highlighting these overlaps and regulatory interactions were generated using the plotBrowserTrack() function from ArchR.

## Data availability

Raw sequencing data, processed Cell Ranger outputs, and comprehensive sample metadata for the human RPE/Choroid cell atlas have been deposited in the Human Cell Atlas (HCA) Data Portal (https://data.humancellatlas.org/hca-bio-networks/eye). The raw sequencing reads for the newly generated datasets have been deposited in the Gene Expression Omnibus (GEO) under accession number GSE312791. Furthermore, raw and normalized count matrices, hierarchical cell-type annotations, and dimensional embeddings are publicly accessible via the CELLxGENE collection (link) and the Cell Annotation Platform (CAP) (https://celltype.info/project/765).

## Code availability

The computational pipeline used for atlas construction and analysis was adapted from the established Human Retina Cell Atlas (HRCA) workflow. All custom scripts and reproducibility environment details are publicly available on GitHub at https://github.com/RCHENLAB/HRCA_reproducibility.

## Acknowledgements

This project was funded by the Chan Zuckerberg Initiative (CZI) grants 2019-002425 and 2021-239847 to R.C.; NIH/NEI R01EY036173 to S.B.; NIH/NIA U54AG075931 to Q.M.; NEI Intramural research Program to K.B.; NIH R01AI180047 and R01EY036519 to D.S; NLM Intramural Research Program (ZIA-LM202401) to R.H.S and A.V.P.; NIH/NEI RO1EY032966 and RO1EY029428 to C.-H.S. Next-generation sequencing (NGS) was performed on instruments supported by the National Institutes of Health (NIH) shared instrument grant S10OD023469 to R.C. This work was also supported by the Pelotonia Institute of Immuno-Oncology (PIIO). The authors acknowledge support for the Gavin Herbert Eye Institute at the University of California, Irvine, from an unrestricted grant from Research to Prevent Blindness, The Paul and Evanina Bell Mackall Foundation Trust and from NIH core grant P30 EY034070. We thank Jennifer Zamanian, Jason Hilton, and the Lattice team at Stanford University for their support with data dissemination. This publication is part of the Human Cell Atlas (http://www.humancellatlas.org/publications/). We also gratefully acknowledge the feedback and discussions from Evan Biederstedt and the members of the HCA Eye Biological Network.

## Author contributions

J.S. and R.C. conceptualized and designed the study. R.C. supervised the project. M.M.D. and J.T.S. provided RPE and Choroid samples. X.B. and Y.L. developed protocols for sample collection and processing. I.Y., X.B., and Y.Z. collected and processed tissue samples. Y.L. supervised and conducted the snRNA-seq and snATAC-seq experiments. Y.Z. conducted the Xenium experiments. J.W., Ji.L., T.Y., J.J., and H.C. contributed to data analysis. Je.L. performed the benchmarking study of integration methods. S.B., K.B., M.C., D.S., C.-H.S., J.Z., D.O., S.X.Z., R.S., R.H.S., and A.V.P. served as members of the RPE/Choroid Curation Working Group and contributed to the consensus classification of cell types and states. A.G., A.M., and Q.M. performed cell senescence analysis. M.F. and A.C.V. managed data curation and dissemination to the Cell Type Annotation Platform. J.S. wrote the initial draft of the manuscript. All authors reviewed the manuscript and contributed to its critical revision.

