## Supplemental figures for "An Integrated Multi-omics Single Cell Atlas of the Human RPE and Choroid"

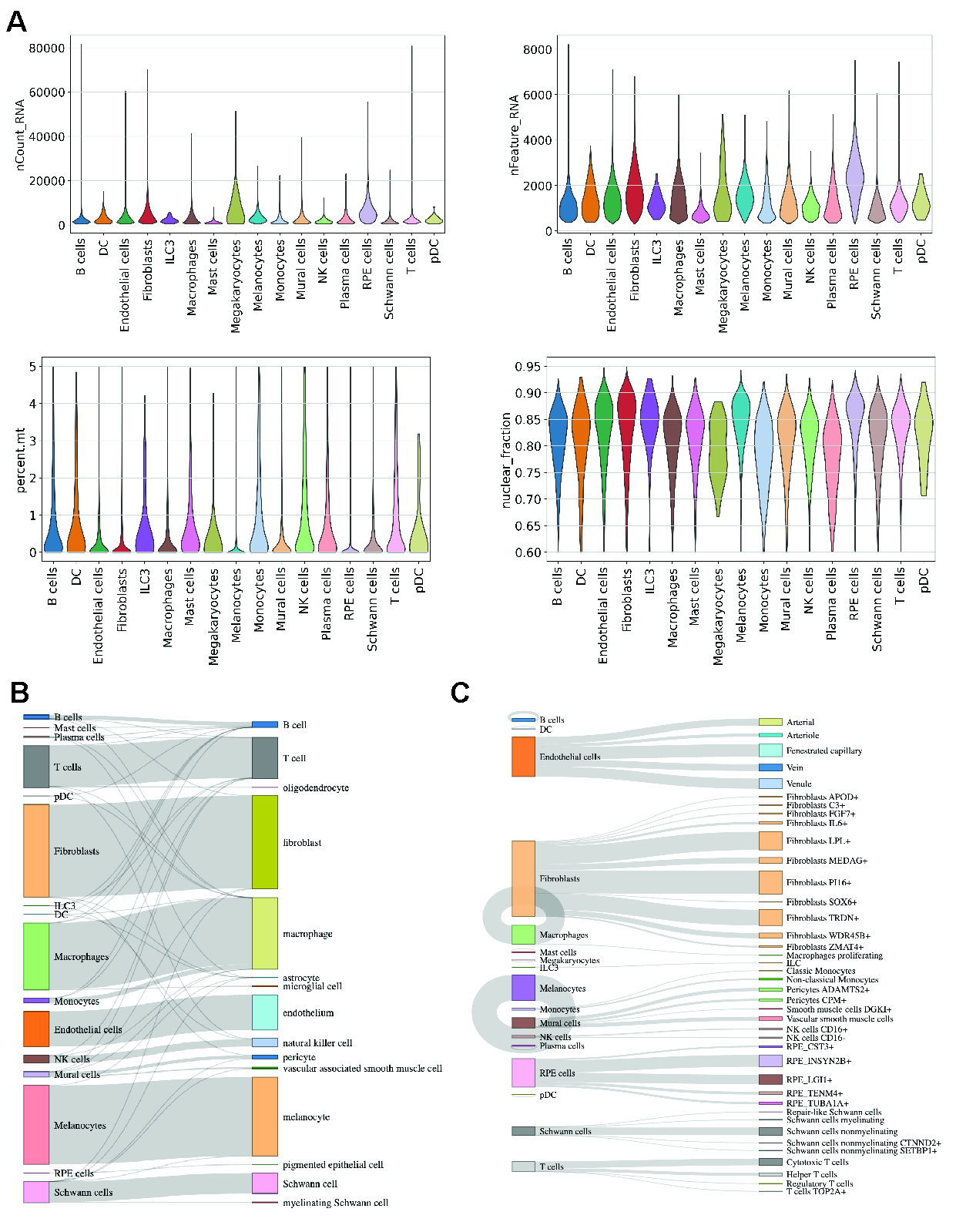

**Figure S1. Quality control and annotation validation of the snRNA-seq atlas.**

**a,** Violin plots displaying quality control metrics, including total UMI counts, number of detected genes, percentage of mitochondrial reads, and the nuclear fraction per nucleus. **b,** Sankey diagram illustrating the high consistency between the cell-type annotations in this study and the metadata labels from a previously published landmark study. **c,** Hierarchical overview of cell-type composition in the snRNA-seq dataset at both major-class and subclass (48 states) resolutions.

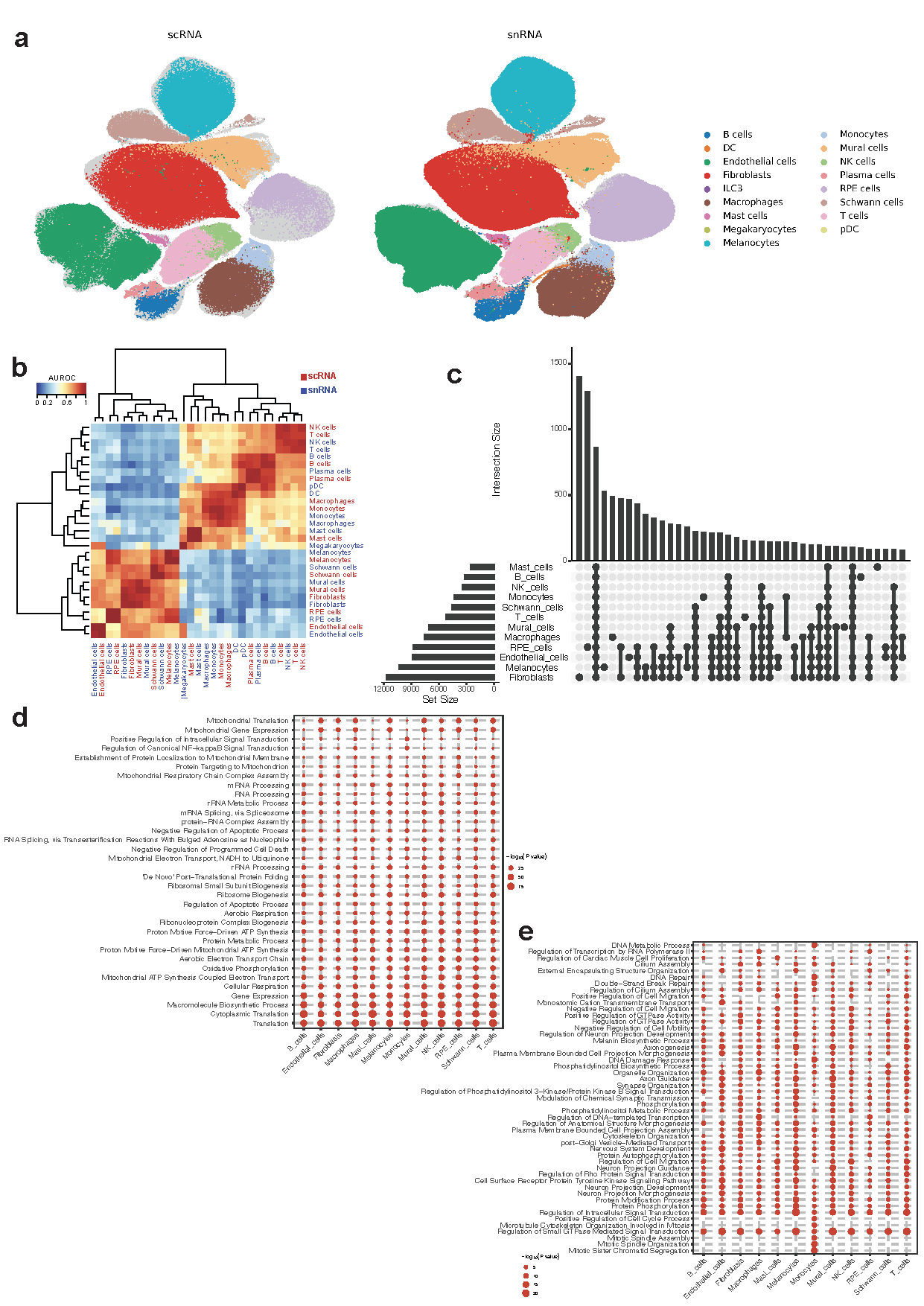

**Figure S2. Modality-driven transcriptomic similarities and differences in the human RPE/choroid.**

**a,** UMAP visualization demonstrating the co-embedding of scRNA-seq and snRNA-seq datasets using sysVI. **b,** Heatmap quantifying transcriptomic similarity between snRNA-seq (blue) and scRNA-seq (red) major classes; color scale represents mean AUROC values derived from MetaNeighbor self-projection analyses. **c,**UpSet plot summarizing the intersection of modality-specific differentially detected genes (DDGs) across major cell classes. **d-e,** Gene Ontology (GO) enrichment analysis of biological processes for genes preferentially detected in scRNA-seq **d** and snRNA-seq **e** Red dots indicate significantly enriched terms (FDR < 0.05).

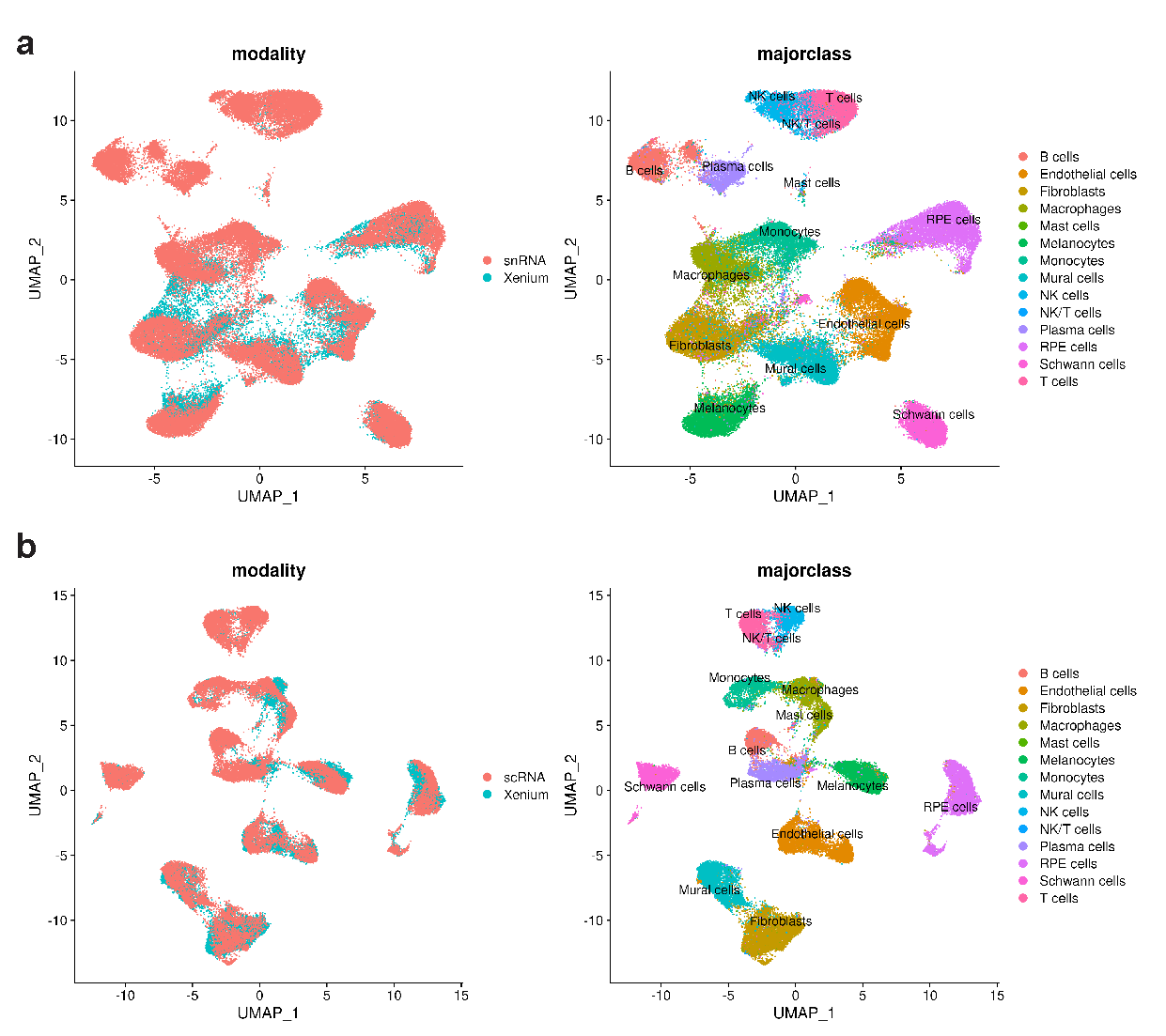

**Figure S3. Integrated co-embedding of Xenium spatial transcriptomics with sequencing modalities.**

**a,** UMAP visualization showing the co-embedding of Xenium spatial data with the snRNA-seq atlas. **b,** UMAP visualization showing the co-embedding of Xenium spatial data with the scRNA-seq atlas.

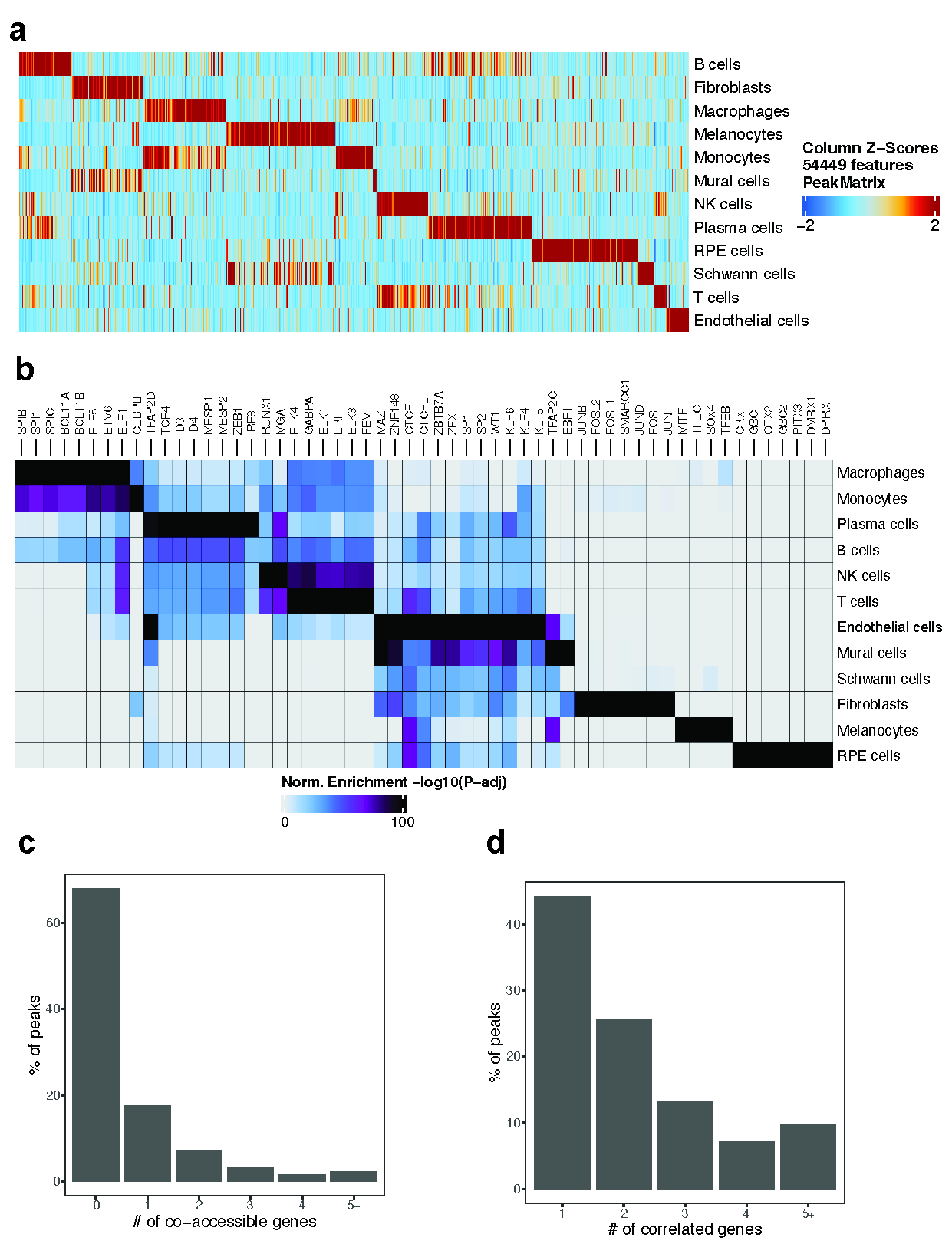

**Figure S4. Cell-type-specific chromatin accessibility, transcription factor enrichment, and distal regulatory linkages.**

**a,** Heatmap displaying marker peaks across major RPE and choroid cell types. Each column represents a specific marker peak. **b,** Heatmap of enriched transcription factor (TF) binding motifs identified within the marker peaks of each cell type. **c,** Distribution of the number of genes exhibiting a co-accessible promoter for each scATAC-seq peak. **d,** Distribution of the number of genes with significantly correlated expression for each scATAC-seq peak.

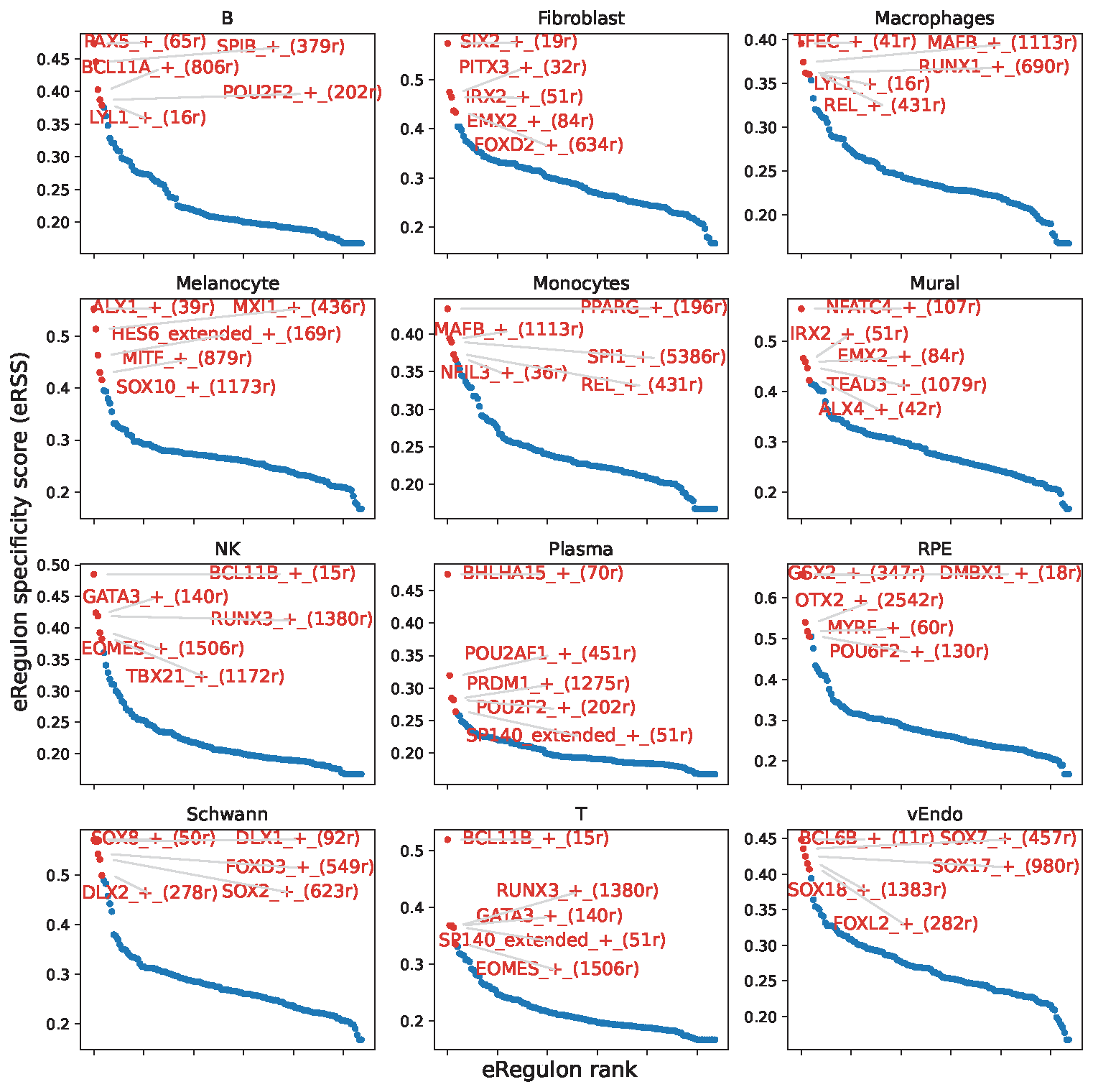

**Figure S5. Ranked eRegulon Specificity Score (eRSS) plots for major cell types in the RPE and choroid atlas.**

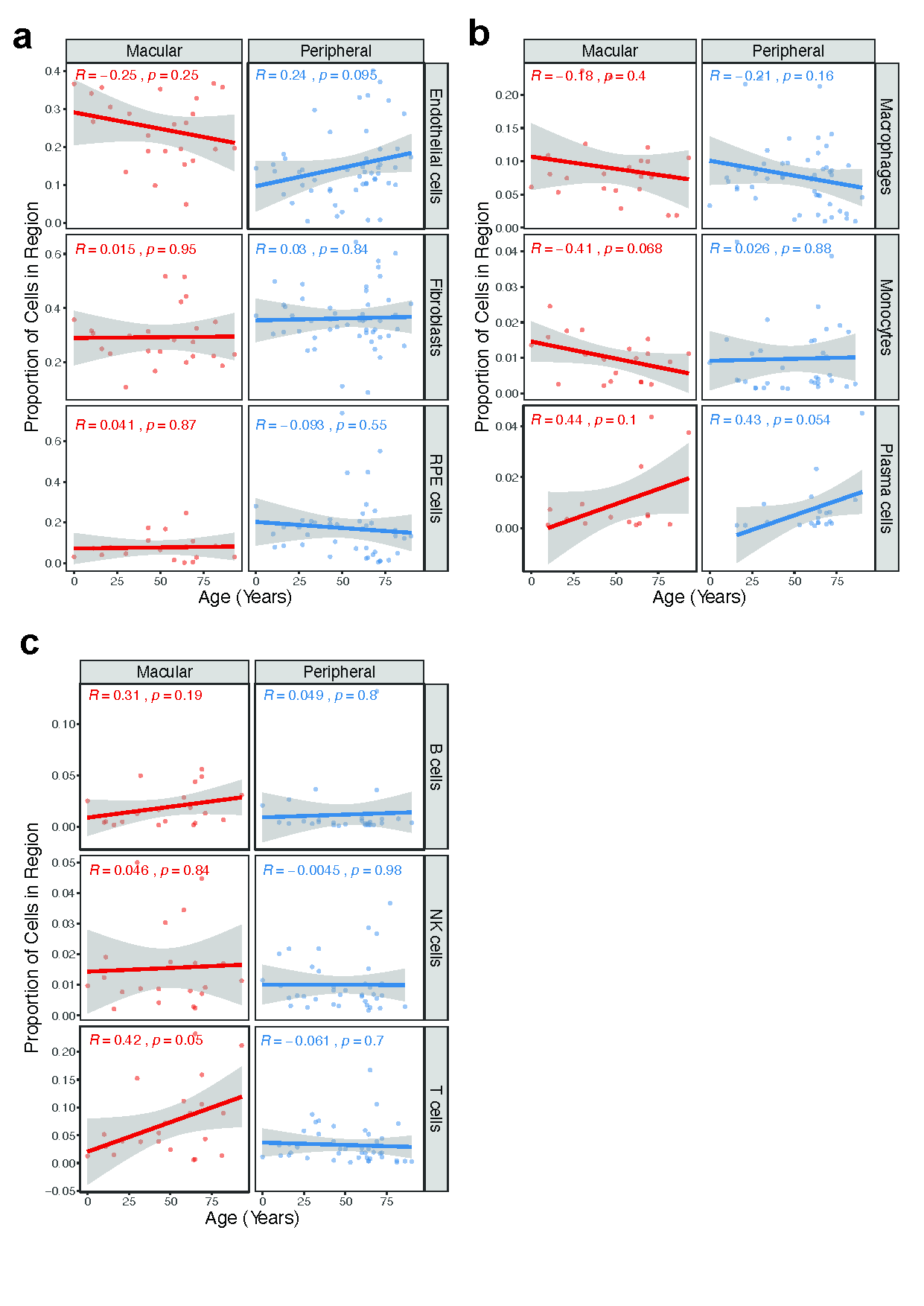

**Figure S6.** Age-associated shifts in cellular composition for macular and peripheral **a,** endothelial cells, fibroblasts, RPEs, **b,** macrophages, monocytes, plasma cells, **c,** B cells, NK cells, and T cells.

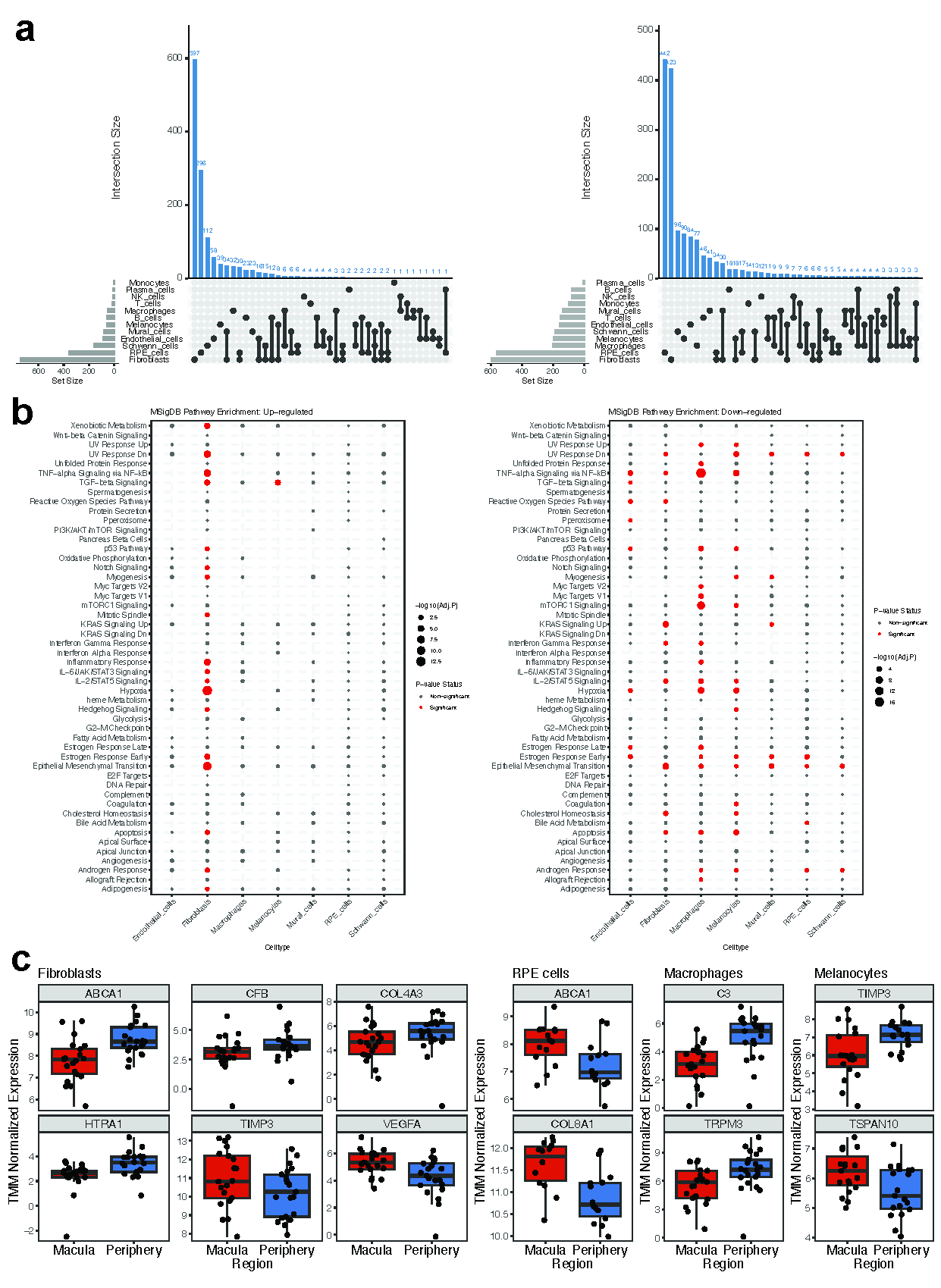

**Figure S7. Transcriptomic divergence between macular and peripheral cell populations.**

**a,** UpSet plot illustrating regional DEGs that are unique to specific cell types or shared across multiple lineages. **b,** Bubble plot displaying MSigDB pathways enriched in macular-upregulated genes (left panel) and peripheral-upregulated genes (right panel). **c,** Box plots showing TMM-normalized expression levels of regional DEGs that intersect with known AMD risk genes in fibroblasts, RPE cells, macrophages, and melanocytes (left to right).

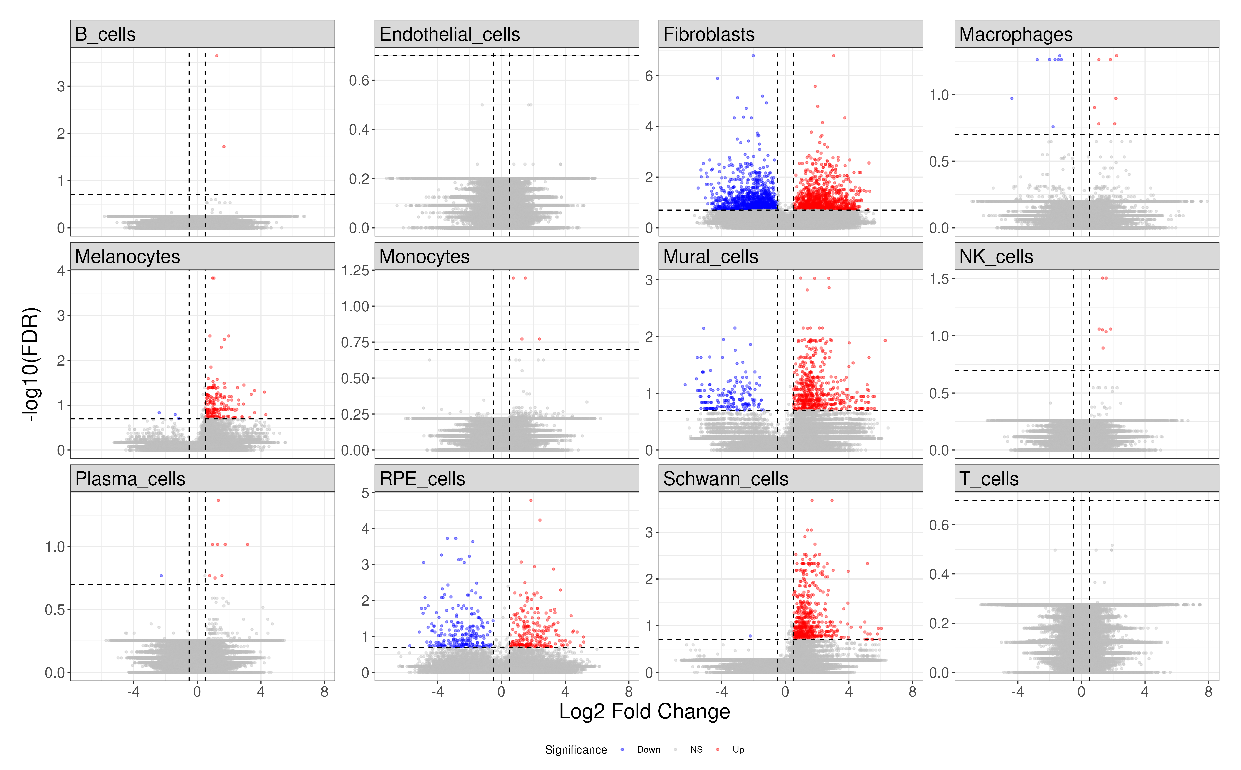

**Figure S8. Differential chromatin accessibility across 12 major cell types.**

Volcano plots illustrating regional DARs for each cell type. Red dots indicate peaks significantly enriched in macular cells (FDR ≤ 0.2, Log_2FC ≥ 0.5), while blue dots indicate peaks enriched in peripheral cells (FDR ≤ 0.2, Log_2FC ≤ -0.5).

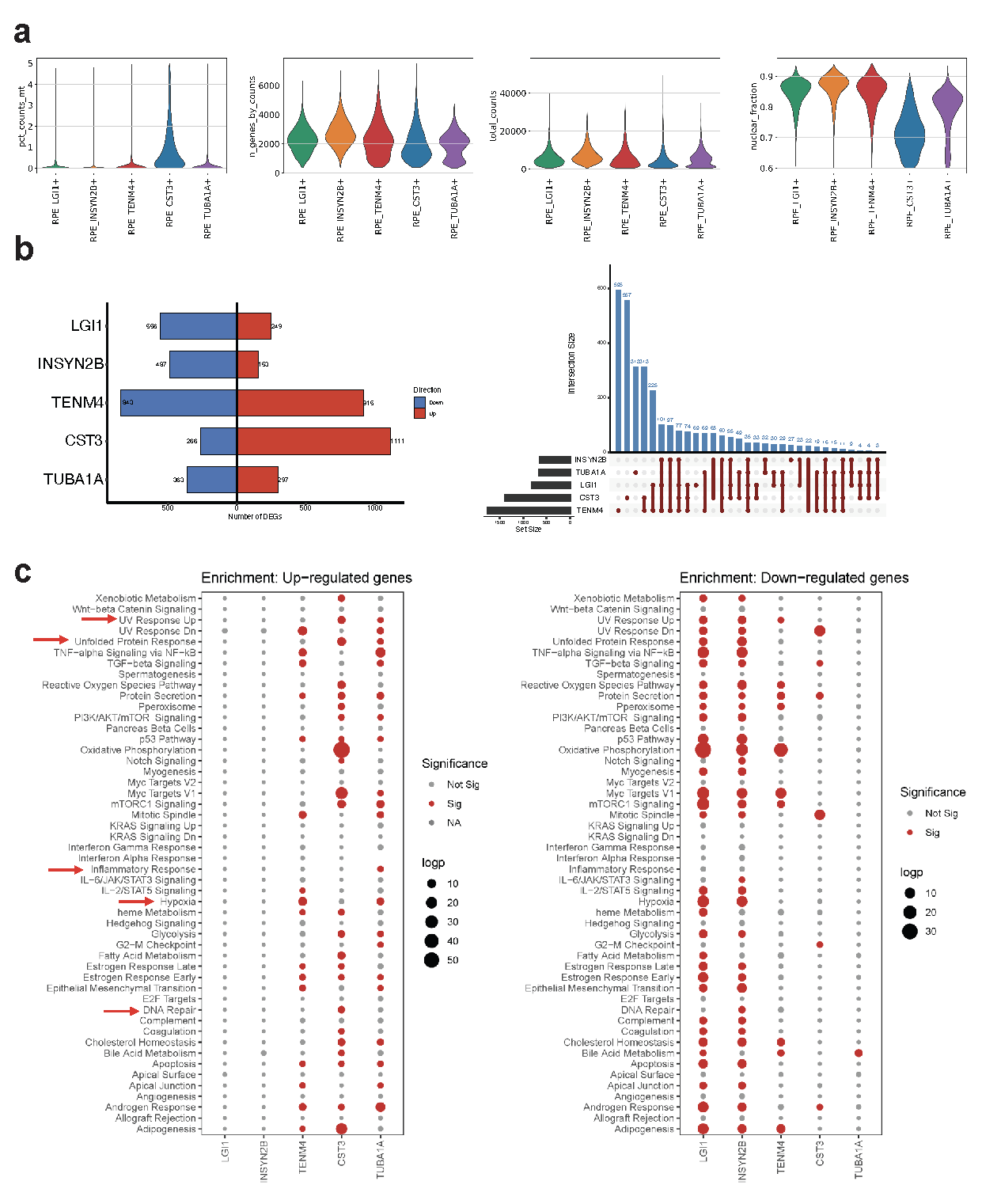

**Figure S9. Quality control metrics and transcriptomic landscape of RPE clusters.**

**a,** Violin plots displaying quality control (QC) metrics across the five defined RPE clusters. Metrics include (from left to right): percentage of mitochondrial transcripts (pct_counts_mt) , number of unique genes detected (n_genes_by_counts) , total UMI counts (total_counts) , and the estimated nuclear fraction. **b,** Differential gene expression analysis across RPE clusters. The bi-directional bar plot (left) quantifies cluster-specific up-regulated (red) and down-regulated (blue) genes. The UpSet plot (right) illustrates the intersection of differentially expressed genes (DEGs), highlighting genes unique to specific clusters or shared across multiple RPE lineages. **c,** Bubble plots summarizing pathway enrichment analysis for up-regulated (left) and down-regulated (right) genes across the RPE clusters. The size and color of the bubbles correspond to the significance of enrichment (-log_10_FDR).

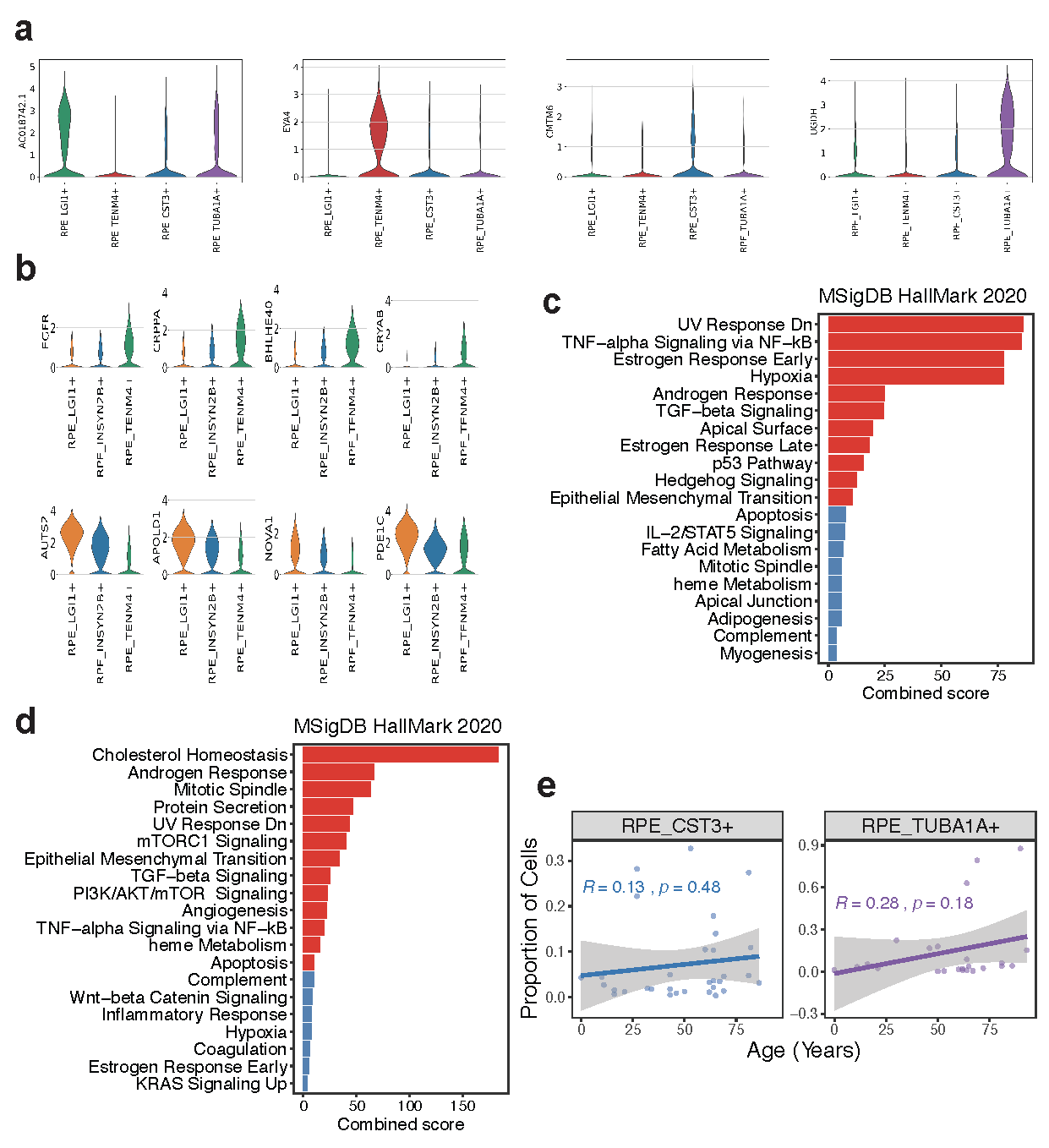

**Figure S10. Transcriptomic markers and functional pathways of the RPE spatial continuum.**

**a,** Violin plots displaying the expression of representative marker genes in the snRNA-seq atlas used to annotate RPE subtypes within the Xenium spatial transcriptomics data. **b,** RNA velocity-derived pseudotime analysis identifying genes positively (top) and negatively (bottom) associated with the central-to-peripheral spatial transition. Violin plots show the expression of these genes across the three RPE subtypes that constitute the RPE spatial continuum. **c,** Bar plot illustrating MSigDB Hallmark pathway enrichment for genes positively correlated with the peripheral transition. **d,** Bar plot illustrating MSigDB Hallmark pathway enrichment for genes correlated with central RPE localization. **e,** Scatter plots and linear regression analysis depicting age-associated shifts in the cellular proportions of CST3+ and TUBA1A+ RPE cells.

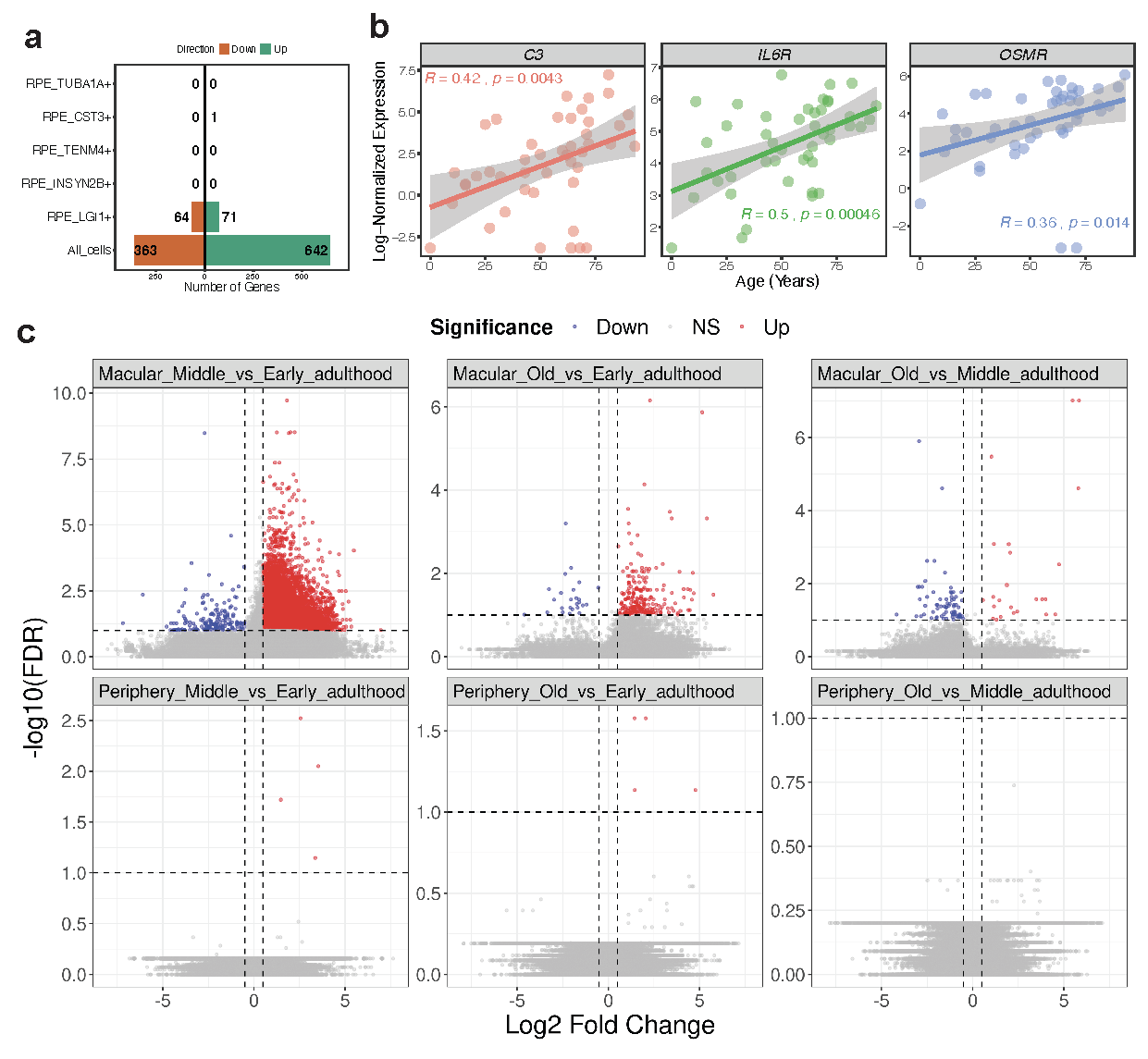

**Figure S11. Regional and temporal transcriptomic shifts in the aging RPE.**

**a,** Bi-directional bar plot quantifying the number of aging-DEGs identified across specific RPE clusters and in the pan-RPE pool ("All_cells"). Red indicates up-regulated genes, and blue indicates down-regulated genes. **b,** Scatter plots illustrating the positive correlation between donor age and the log-normalized expression of inflammatory markers C3, IL6R, and OSMR. **c,** Volcano plots displaying aging-DEGs across three life-stage transitions (Middle vs. Early, Old vs. Early, and Old vs. Middle adulthood) in both macular and peripheral RPE. Red and blue dots denote significantly up-regulated and down-regulated peaks, respectively.

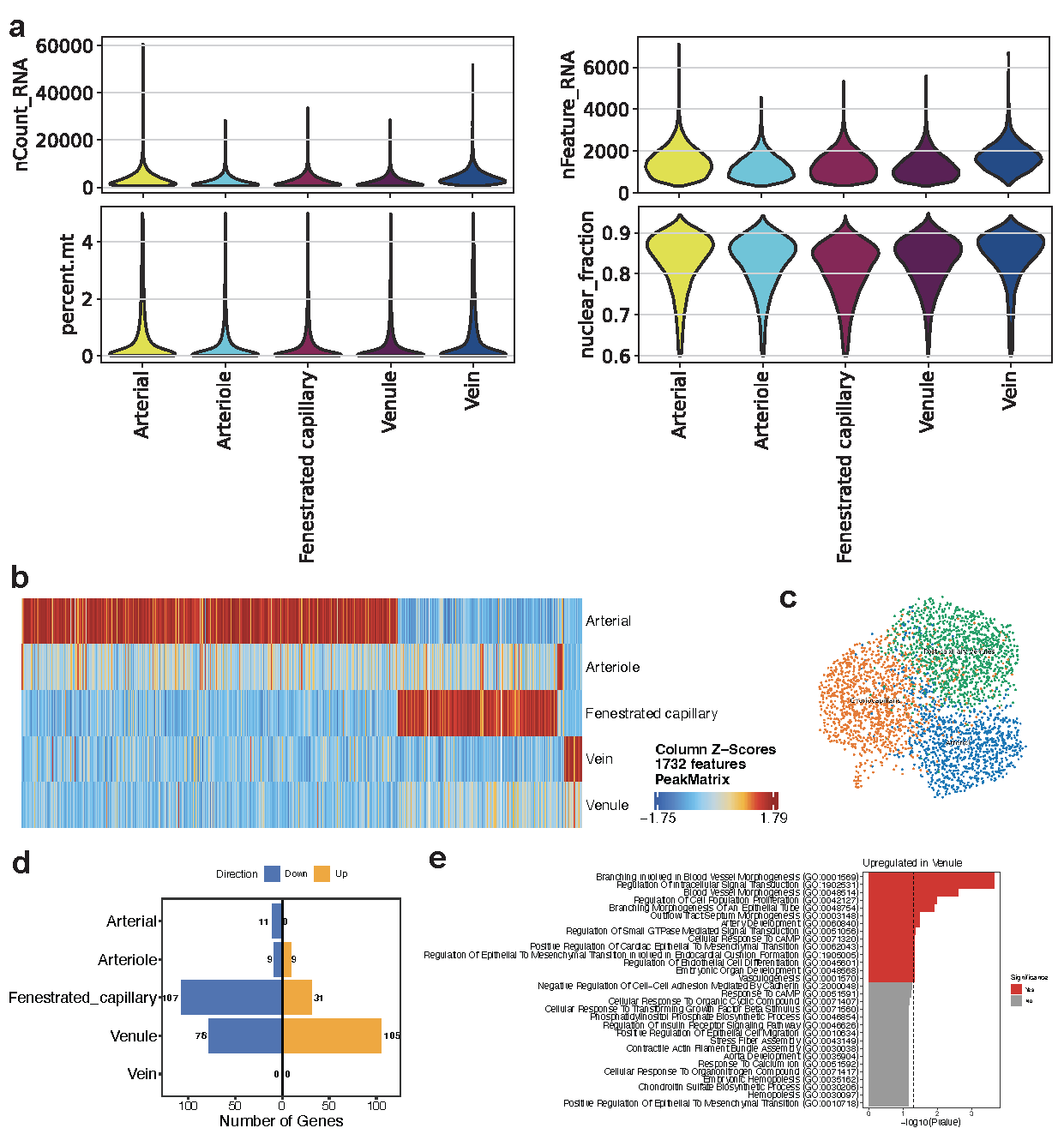

**Figure S12. Quality control and transcriptomic/epigenetic heterogeneity of choroidal endothelial cell subtypes.**

**a,** Violin plots displaying quality control (QC) metrics across the five defined endothelial cell (EC) subtypes: arterial, arteriole, fenestrated capillary, venule, and vein. Metrics include (from top-left to bottom-right): total UMI counts, number of unique genes detected, percentage of mitochondrial transcripts, and the estimated nuclear fraction. **b,** Heatmap illustrating the Z-score normalized accessibility of 1,732 marker peaks across the EC subtypes. **c,** UMAP visualization of endothelial cells based on Xenium spatial transcriptomics dat. **d,** Bi-directional bar plot quantifying the number of differentially expressed genes (DEGs) in the macula compared to the periphery across the five EC subtypes. Red bars indicate up-regulated genes, and blue bars indicate down-regulated genes. **e,** Bar plot showing Gene Ontology (GO) pathway enrichment for genes significantly up-regulated in macular venule ECs.

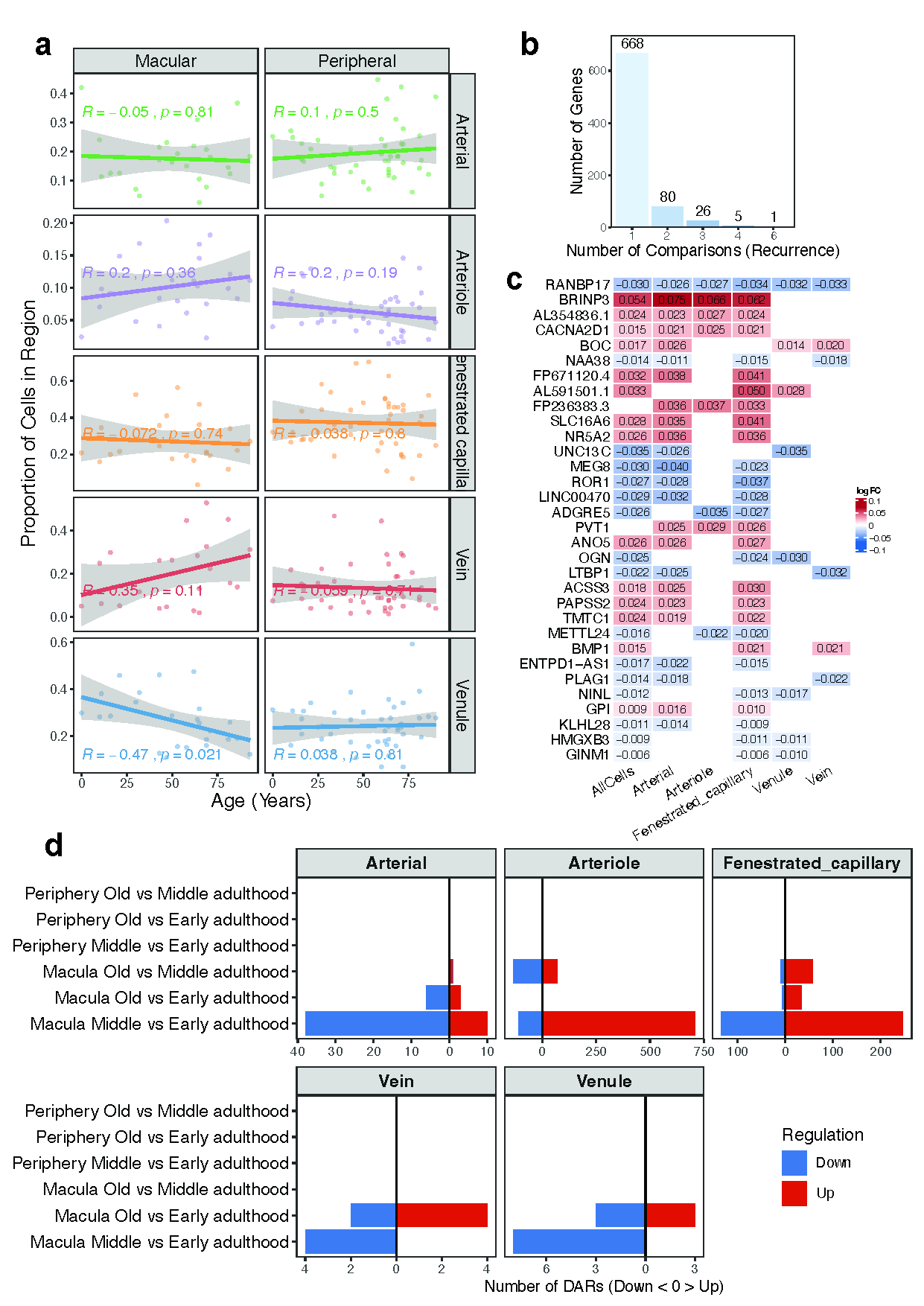

**Figure S13. Age-associated regional and cellular shifts in choroidal endothelial cells.**

**a,** Scatter plots and linear regression analysis depicting age-associated shifts in the cellular proportions of five endothelial cell (EC) subtypes, stratified by region (macular vs. peripheral). **b,** Bar plot illustrating the recurrence of age-associated differentially expressed genes (aging-DEGs) across EC subtypes. The x-axis indicates the number of comparisons (recurrence) in which a gene was identified as significantly differentially expressed. **c,** Heatmap displaying the aging effect (expressed as logFC) for selected genes across all ECs combined ("AllCells") and within the five individual EC subtypes. **d,** Bi-directional bar plot quantifying the number of aging-DARs identified across the five EC subtypes. Comparisons represent three life-stage transitions (Middle vs. Early, Old vs. Early, and Old vs. Middle adulthood) in both macular and peripheral regions. Red and blue bars denote up-regulated and down-regulated accessibility, respectively.

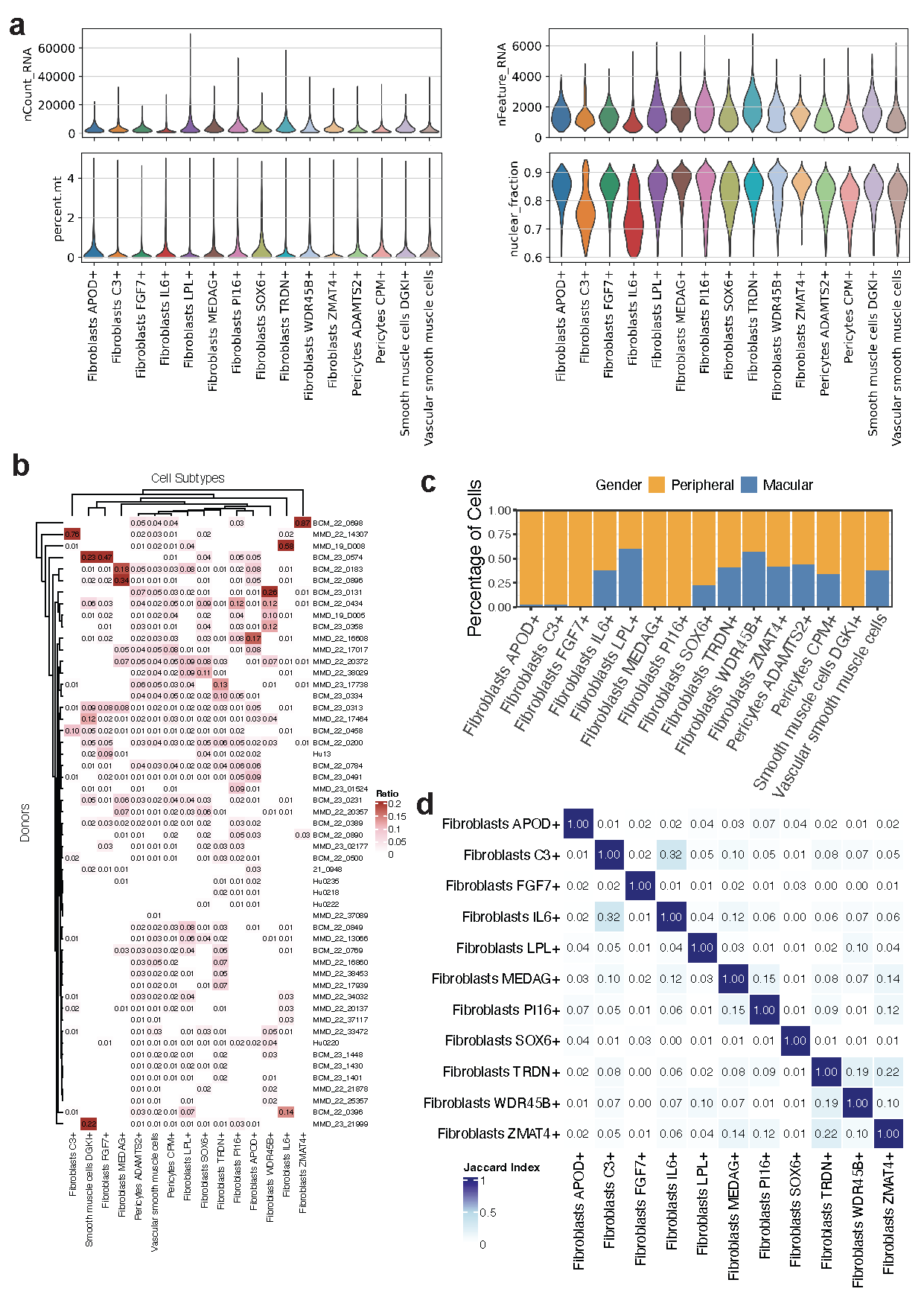

**Figure S14. Quality control and distribution of choroidal stromal cell subtypes.**

**a,** Violin plots displaying quality control (QC) metrics across the defined stromal, pericyte, and smooth muscle cell clusters. Metrics include (from top-left to bottom-right): total UMI counts, number of unique genes detected, percentage of mitochondrial transcripts, and the estimated nuclear fraction. **b,** Heatmap illustrating the percentage of cells contributed by each profiled donor across the stromal and perivascular sub-clusters. **c,** Bar plot showing the regional distribution (macula vs. periphery) of cells across all defined stromal and perivascular types and states. **d,** Heatmap displaying the pairwise Jaccard index across the defined stromal types and states, indicating the degree of transcriptomic similarity or overlap between the identified clusters.

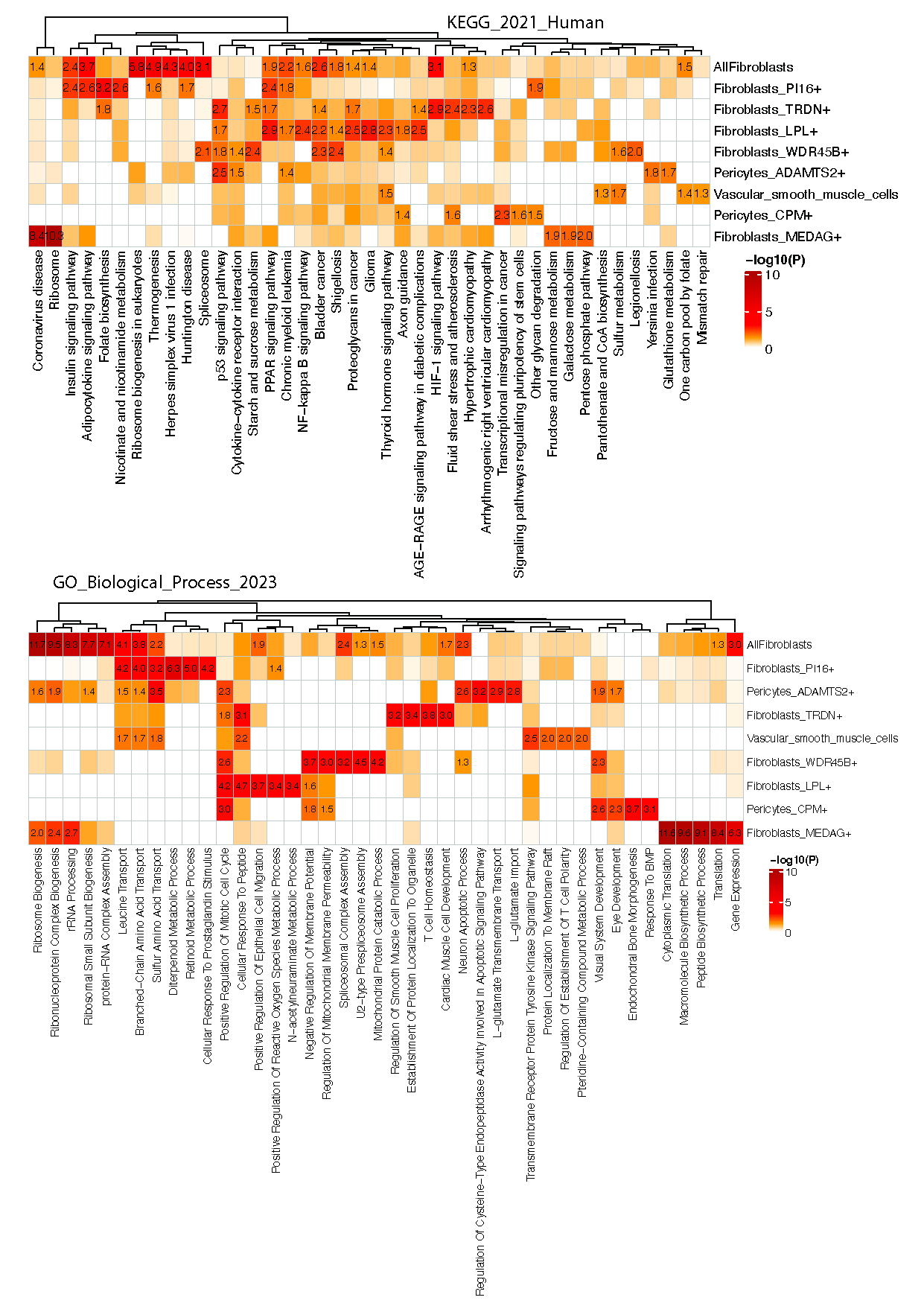

**Figure S15. Pathway and Gene Ontology enrichment of age-associated differentially expressed genes in choroidal stromal cells.** Heatmap illustrating enriched biological pathways and Gene Ontology (GO) terms for genes significantly regulated with age (aging-DEGs) across all fibroblasts ("AllFibroblasts") and specific stromal and perivascular subtypes.

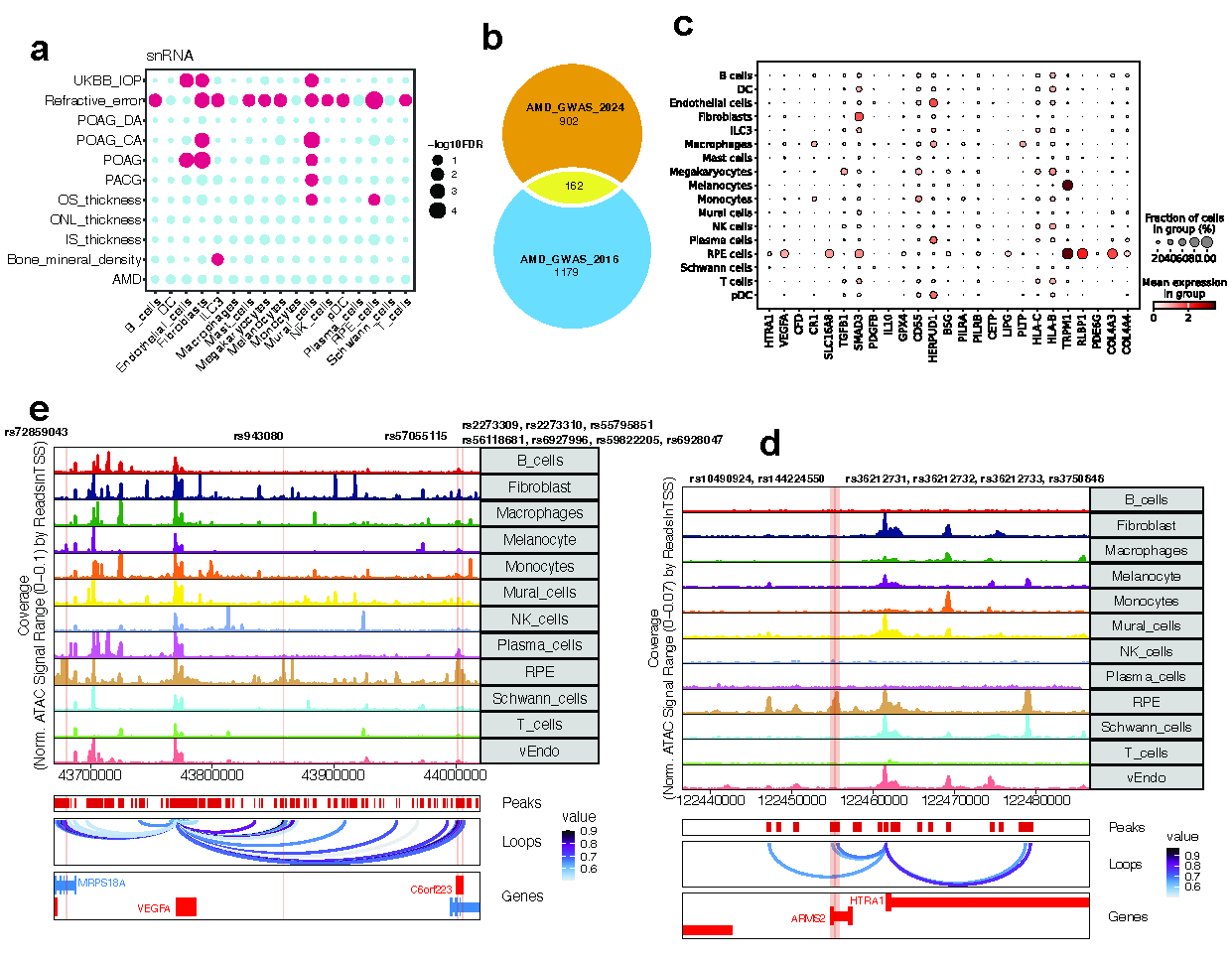

**Figure S16. Integration of AMD GWAS loci with multi-omic RPE/Choroid cell-type landscapes.**

**a,** Cell-type enrichment analysis of various GWAS traits based on gene expression profiles across major RPE and choroidal cell classes. b, Venn diagram illustrating the overlap of 95% credible set variants from two landmark AMD GWAS studies. **c,** Dot plot showing the expression patterns of prioritized AMD candidate genes across major cell types from snRNA-seq. **d,** Genome track visualization of the ARMS2/HTRA1 risk locus. Highlighted regions (firebrick) indicate open chromatin regions (OCRs) harboring AMD GWAS variants that are linked to HTRA1 via peak-to-gene correlations. **e,** Genome track visualization of the VEGFA locus. Highlighted regions (firebrick) denote OCRs and their associated AMD GWAS variants that are correlated with VEGFA expression in specific cell types.

**Supplemental Tables:**

Supplementary Table S1. Metadata for scRNA-seq and snRNA-seq samples

Supplementary Table S2. Donor contribution to each major cell class for snRNA-seq

Supplementary Table S3. NS-Forest markers of major cell classes for snRNA-seq

Supplementary Table S4. Modality-specific differentially detected genes per major cell class

Supplementary Table S5. Xenium panel gene list

Supplementary Table S6. Metadata for snATAC-seq samples

Supplementary Table S7. Donor contribution to each major cell class for snATAC-seq

Supplementary Table S8. Differentially accessible regions per major cell class for snATAC-seq

Supplementary Table S9. List of co-accessible regions

Supplementary Table S10. List of peak-to-gene links

Supplementary Table S11. Regional differentially expressed genes per major cell class for snRNA-seq

Supplementary Table S12. Comparison of regional DEGs between this study and Voigt et al. (PNAS, 2019)

Supplementary Table S13. Regional differentially accessible regions per major cell class for snATAC-seq

Supplementary Table S14. Donor contributions to RPE types and states

Supplementary Table S15. NS-Forest markers for RPE types and states

Supplementary Table S16. Cluster-specific differentially expressed genes of RPE types and states

Supplementary Table S17. RNA velocity pseudotime-associated genes of RPE types

Supplementary Table S18. Aging-associated differentially expressed genes of pan-RPE cells

Supplementary Table S19. Regional differentially expressed genes per endothelial cell type

Supplementary Table S20. Regional differentially accessible regions per endothelial cell type

Supplementary Table S21. Aging-associated differentially expressed genes of endothelial cells

Supplementary Table S22. Differential chromatin accessibility regions in macular tissue: middle vs. early adulthood

Supplementary Table S23. Cluster-specific differentially expressed genes of stromal types and states

Supplementary Table S24. Aging-associated differentially expressed genes of stromal cells

Supplementary Table S25. Functional prioritization of AMD GWAS risk variants
